# Flagella Ca^2+^ elevations regulate pausing of retrograde intraflagellar transport trains in adherent *Chlamydomonas* flagella

**DOI:** 10.1101/2020.08.17.240366

**Authors:** Cecile Fort, Peter Collingridge, Colin Brownlee, Glen Wheeler

## Abstract

The movement of ciliary membrane proteins is directed by transient interactions with intraflagellar transport (IFT) trains. The green alga *Chlamydomonas* has adapted this process for gliding motility, using IFT to move adhesive glycoproteins (FMG-1B) in the flagella membrane. Although Ca^2+^ signalling contributes directly to the gliding process, uncertainty remains over the mechanisms through which Ca^2+^ acts to influence the movement of IFT trains. Here we show that flagella Ca^2+^ elevations regulate IFT primarily by initiating the movement of paused retrograde IFT trains. Flagella Ca^2+^ elevations exhibit complex spatial and temporal properties, including high frequency repetitive Ca^2+^ elevations that prevent the accumulation of paused retrograde IFT trains. We show that flagella Ca^2+^ elevations disrupt the IFT-dependent movement of microspheres along the flagella membrane. The results suggest that flagella Ca^2+^ elevations directly disrupt the interaction between retrograde IFT particles and flagella membrane glycoproteins to modulate gliding motility and the adhesion of the flagellum to a surface.

## Introduction

Cilia and flagella are microtubule-rich organelles that extend from the surface of many eukaryote cell types. Cilia play well-characterised roles in motility, but also have additional roles as important cellular sensors. The elongated morphology of eukaryote cilia (length 1-20 µm, diameter 200 nm) requires specialised mechanisms to transport proteins along its length. This is achieved through the process of intraflagellar transport (IFT), in which cargo proteins required for the assembly, maintenance and sensory functions of cilia are attached to large protein complexes (IFT particles) and moved along the axoneme through the activity of microtubule motors^1^. Kinesin-2 directs movement away from the cell body (anterograde), while dynein-1b (dynein 2 in mammals) mediates the return of IFT particles towards the cell (retrograde). The anterograde IFT particles are assembled into trains, which move in a processive manner to the ciliary tip. Once the anterograde IFT particles reach the ciliary tip, kinesin dissociates from the IFT particle and returns to the cell body via diffusion. The diffusive return of kinesin to the cell body may play a critical role in regulating ciliary length, acting to limit the rate at which IFT particles can enter the flagellum ^2, 3^. IFT trains at the ciliary tip dissociate into multiple smaller trains and move back towards the cell body ^2^. Although dynein-1b is present on anterograde IFT particles, it is carried along the axoneme in an auto-inhibited form until it reaches the flagella tip to prevent a tug-of-war between the two microtubule motors ^4^.

The import, export and movement of many ciliary proteins is determined by their interactions with IFT particles. IFT particles are composed of around 20 proteins arranged into two sub-complexes (IFT-A and IFT-B). Proteins within the IFT-B complex have specific roles in binding to cargo proteins including important structural components such as such as tubulin (IFT74/81) and outer arm dynein (IFT46) ^5, 6^. IFT-A proteins interact with cargo proteins involved in signalling pathways, such as G-protein coupled receptors^7^. Expression of a truncated IFT-A protein (IFT140) in *Chlamydomonas* flagella resulted in a reduced complement of membrane-associated proteins, such as small GTPases and ion channels ^8^. An additional protein complex, known as the BBsome, acts as a cargo adapter and is implicated in the transport of a range of membrane-associated ciliary proteins ^9^. Distinct proteins within the IFT complex play therefore specific roles in binding to cargo proteins, although the factors regulating interactions between cargo proteins and IFT particles are less well-characterised. Improved knowledge of these processes is required to better understand the mechanisms determining the movement and distribution of ciliary proteins.

There is evidence that loading and unloading of some ciliary proteins is a highly regulated process ^10^. Second messengers such as Ca^2+^ and cAMP may also contribute to the regulation of cargo binding ^11^. For example, mice lacking the ciliary localised Ca^2+^-permeable ion channel PKD2L1 exhibit defects in the IFT-dependent trafficking of the Shh transcription factor Gli2 to the tip of the primary cilium ^12^. In *Chlamydomonas*, phosphorylation of the FLA8 kinesin subunit by the Ca^2+^-dependent protein kinase CDPK1 blocks FLA8 entry in to the flagellum and promotes dissociation of kinesin from the IFT particles at the ciliary tip, suggesting a role for Ca^2+^ in cargo unloading ^13^.

Calcium has also been implicated in the movement of flagella membrane proteins in *Chlamydomonas*. An adhesive glycoprotein in the flagella membrane (FMG-1B) allows *Chlamydomonas* cells to adhere to substrates ^14-16^. Transient interactions between FMG-1B and IFT particles direct the movement of FMG-1B along the flagella, demonstrated by the co-localisation of individual IFT trains with polystyrene microspheres moving along the flagella surface ^17^. This process is utilised to drive gliding motility in *Chlamydomonas*, where the cell moves along a surface on its adherent flagella. The motile force is provided by the retrograde movement of FMG-1B along the flagellum, which results in the flagellum pulling the cell body forward ^15, 18^. Gliding motility is therefore caused by transient associations between FMG-1B and retrograde IFT particles, representing an excellent system in which to study dynamic interactions between cargo proteins and IFT particles.

Gliding motility is regulated by Ca^2+^, requiring micromolar Ca^2+^ in the external media ^19^. Bead movement is also inhibited by the absence of Ca^2+^ or by the presence of diltiazem, an inhibitor of voltage-gated Ca^2+^ channels ^19^. The implication of IFT motors in gliding suggests that Ca^2+^ most likely acts to regulate the interaction between IFT and FMG-1B. Shih *et al* ^17^ found that chelating external Ca^2+^ reduced the pausing frequency of IFT trains, suggesting that Ca^2+^ was required for the interaction between FMG-1B and IFT particles. In contrast, Collingridge et al ^20^ found that the absence of external Ca^2+^ led to a substantial accumulation of IFT particles in adherent flagella, suggesting that Ca^2+^ acts to disrupt rather than promote the interaction between FMG-1B and the IFT particle. This was supported by simultaneous imaging of intraflagellar Ca^2+^ ([Ca^2+^]_fla_) and IFT using biolistically-loaded dextran-conjugated Ca^2+^-responsive fluorescent dyes, which showed that [Ca^2+^]_fla_ elevations acted to initiate the movement of paused IFT particles ^20^.

The discrepancy between these two findings cannot easily be resolved, yet it clearly indicates that greater understanding of the interaction between [Ca^2+^]_fla_ and IFT is required. In particular, the nature of paused anterograde and retrograde IFT trains, and the role of [Ca^2+^]_fla_ in regulating this process, requires more detailed study. Direct evidence for a regulatory function for [Ca^2+^]_fla_ in the interaction between FMG-1B and the IFT particle is currently lacking. The interaction between FMG-1B and IFT may also influence other aspects of flagella function, such as flagella adhesion. Recent observations indicate that the adhesive properties of *Chlamydomonas* flagella are highly dynamic, with blue light promoting rapid changes in flagella adhesion to a surface ^21, 22^. Light-dependent adhesion appears to involve a redistribution of FMG-1B along the flagella surface ^22^, which may therefore be dependent on interactions with IFT. Ca^2+^-dependent signalling processes also contribute to many other flagellar functions in *Chlamydomonas*, such as changes in flagellar beat and waveform during motile responses, flagellar adhesion during mating, flagellar excision and flagellar length control ^23-26^. A more detailed study of the nature of Ca^2+^ signalling in flagella is required to help us understand how Ca^2+^ acts in these multiple roles within flagella and how specificity for each signalling role is achieved.

In this study, we have examined the interactions between [Ca^2+^]_fla_ and IFT in adherent *Chlamydomonas* flagella. Using genetically encoded Ca^2+^ reporters targeted to the flagella, we demonstrate that intraflagellar Ca^2+^ elevations act primarily to regulate the accumulation of paused retrograde IFT particles, by promoting their dissociation from the flagella membrane. We also reveal complex spatiotemporal characteristics of Ca^2+^ in *Chlamydomonas* flagella, such as repetitive high frequency Ca^2+^ spiking and the presence of localised or propagating [Ca^2+^]_fla_ elevations, showing that Ca^2+^ acts locally in its interaction with IFT. By regulating the interaction between IFT and FMG-1B, [Ca^2+^]_fla_ elevations play an important role in modulating flagella adhesion.

## Results

### Distinct forms of paused IFT particles in *Chlamydomonas* flagella

We have previously observed that accumulations of paused IFT trains are associated with gliding motility in adherent flagella ^20^. To examine the nature of the paused IFT trains in greater detail, we used TIRF microscopy of a reporter strain expressing IFT54-mScarlet (strain IFT54-MS). We observed two clear categories of paused IFT particles (Fig 1A). The first of these consisted of paused anterograde IFT particles, which remain highly localised, and continue in an anterograde direction. Paused anterograde IFT trains may disrupt the movement of other anterograde IFT trains, but do not interfere with retrograde transport ^27^. The second category of paused IFT particles was primarily observed near the distal tip of the flagellum. These accumulations appear more diffuse spatially and characteristically slowly expand in size and fluorescence intensity (Fig 1A). The distal IFT accumulations were periodically cleared by retrograde transport, suggesting that they represent paused retrograde IFT trains. This was supported by the presence of similar distal accumulations of IFT trains in a reporter strain for the dynein light intermediate chain (D1bLIC) of cytoplasmic dynein 1b (the retrograde IFT motor) ^28^ (Fig 1B), whereas similar accumulations were absent in a reporter strain for the KAP subunit of kinesin (the anterograde IFT motor) ^20, 29^. As kinesin returns to the cell body via diffusion and is not present in retrograde IFT trains ^2^, the distal IFT accumulations are likely to be formed by paused retrograde IFT trains.

**Figure 1:**
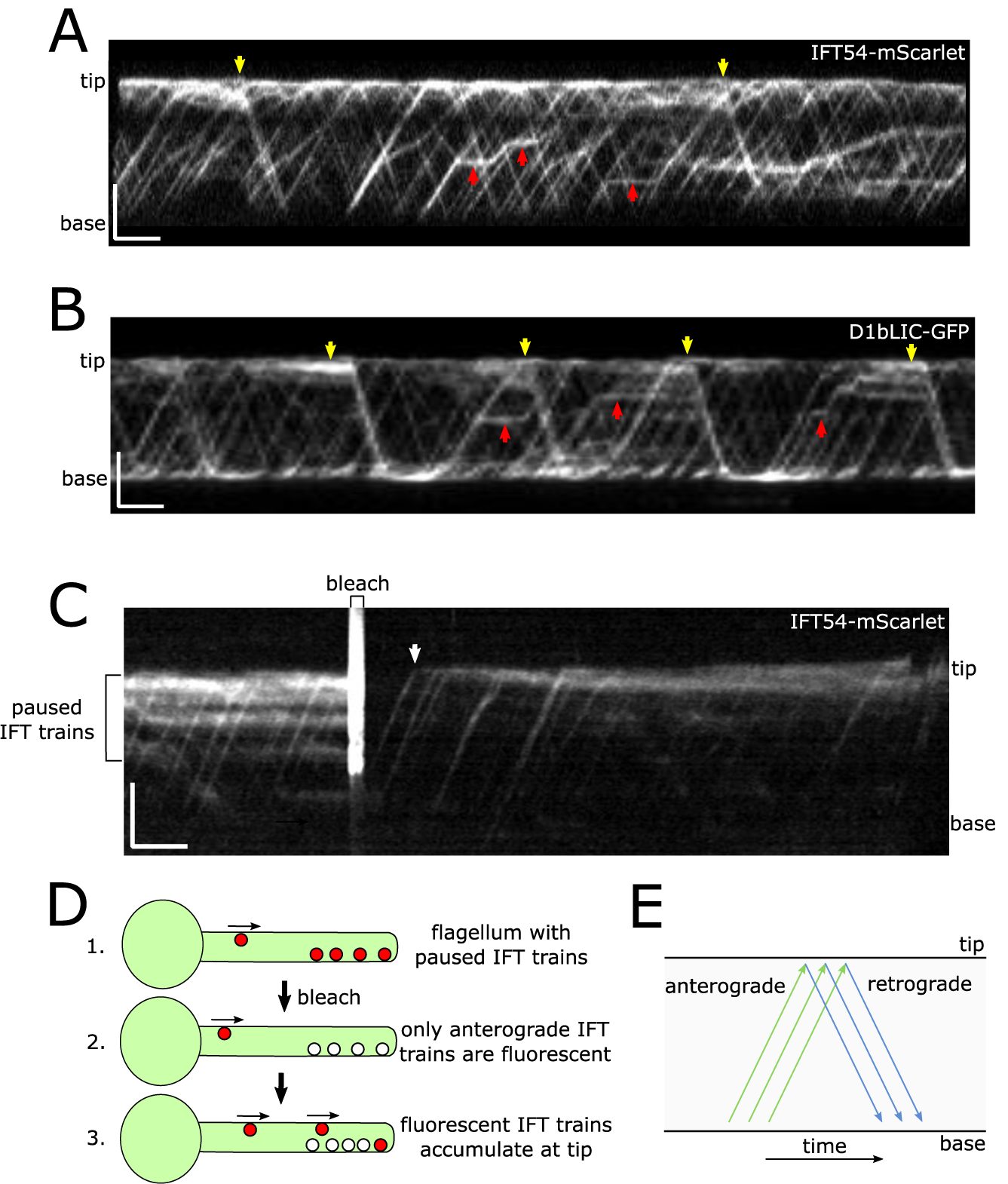
Distinct forms of paused IFT particles in *Chlamydomonas* flagella. **A)** Kymograph of IFT54-mScarlet indicating pausing of IFT trains. Individual anterograde IFT trains can be seen to pause and then continue in an anterograde direction (red arrows). Diffuse accumulations of paused IFT particles in the distal region of the flagellum are periodically cleared in a retrograde direction (yellow arrows). The schematic illustrates the position of paused IFT trains in the kymograph. Bars represent 5 μm and 10 s. **B)** Kymograph of D1bLIC-GFP showing a series of distal accumulations that are cleared by retrograde movement (yellow arrows). **C)** Fluorescence recovery after bleaching (FRAP) of IFT54-mScarlet. IFT54-mScarlet was bleached to allow visualisation of the origin of distal IFT accumulations. After bleaching, IFT54-mScarlet does not begin to accumulate until anterograde IFT trains can be observed to reach the flagella tip (arrow). FRAP experiments were performed on 33 flagella. A representative image is shown. **D**) Schematic representation of FRAP experiment described in (C). **E**) Schematic representation of a kymograph showing the direction of anterograde and retrograde transport.

The diffuse nature of the distal IFT accumulations and their location close to the flagella tip made it difficult to observe directly whether they are composed of retrograde IFT trains. We therefore employed a fluorescence recovery after photobleaching (FRAP) approach to observe how individual IFT trains contribute to these accumulations. Bleaching of flagella exhibiting distal accumulations of paused IFT trains, revealed that individual anterograde IFT trains proceeded to the flagella tip and were largely unimpeded by the accumulation of paused IFT trains (Fig 1C-E). Accumulations of paused IFT trains began to form close to the flagella tip immediately after the arrival of the initial anterograde IFT train after bleaching. The accumulated IFT particles are therefore formed by retrograde IFT particles that pause in the vicinity of the flagella tip immediately after turnaround and departure. The distal IFT accumulations expand in size due to additional retrograde IFT particles arriving at the distal end of the accumulation, rather than either anterograde or retrograde IFT trains adding to the accumulation at its proximal end. As anterograde and retrograde IFT particles travel on distinct microtubules within each doublet ^27^, the accumulation of retrograde IFT particles at the flagella tip does not appear to immediately interfere with anterograde IFT, but may impede the return of retrograde IFT particles.

### Development of a flagella-targeted calcium sensor in *Chlamydomonas*

To identify the role of [Ca^2+^]_fla_ elevations in regulating IFT pausing and other events, we targeted a genetically encoded Ca^2+^-sensitive reporter protein (G-GECO1.1) to *Chlamydomonas* flagella. We fused G-GECO to the C-terminus of CAH6, an abundant carbonic anhydrase that is associated with the flagella membrane and shows an even distribution along the length of the flagellum ^9, 30, 31^ (Fig 2A). Using this approach, we were able to successfully target G-GECO to the flagella of *Chlamydomonas* strain CC5325 (producing strain GG-WT). TIRF microscopy of adherent flagella revealed that [Ca^2+^]_fla_ elevations were associated with gliding movement. [Ca^2+^]_fla_ elevations were observed in 94.8 % of trailing flagella at the onset of the gliding motility (n = 116), but were largely absent from leading flagella (Fig 2B-D; Video 1), strongly supporting previous observations of gliding-related [Ca^2+^]_fla_ elevations using biolistically-loaded Ca^2+^-responsive fluorescent dyes (Oregon Green-BAPTA-dextran) ^20^. Cells expressing CAH6-Venus did not exhibit any changes in fluorescence during gliding motility, indicating that motion artefacts were not responsible for changes in G-GECO fluorescence (Supplementary Fig 1).

**Figure 2:**
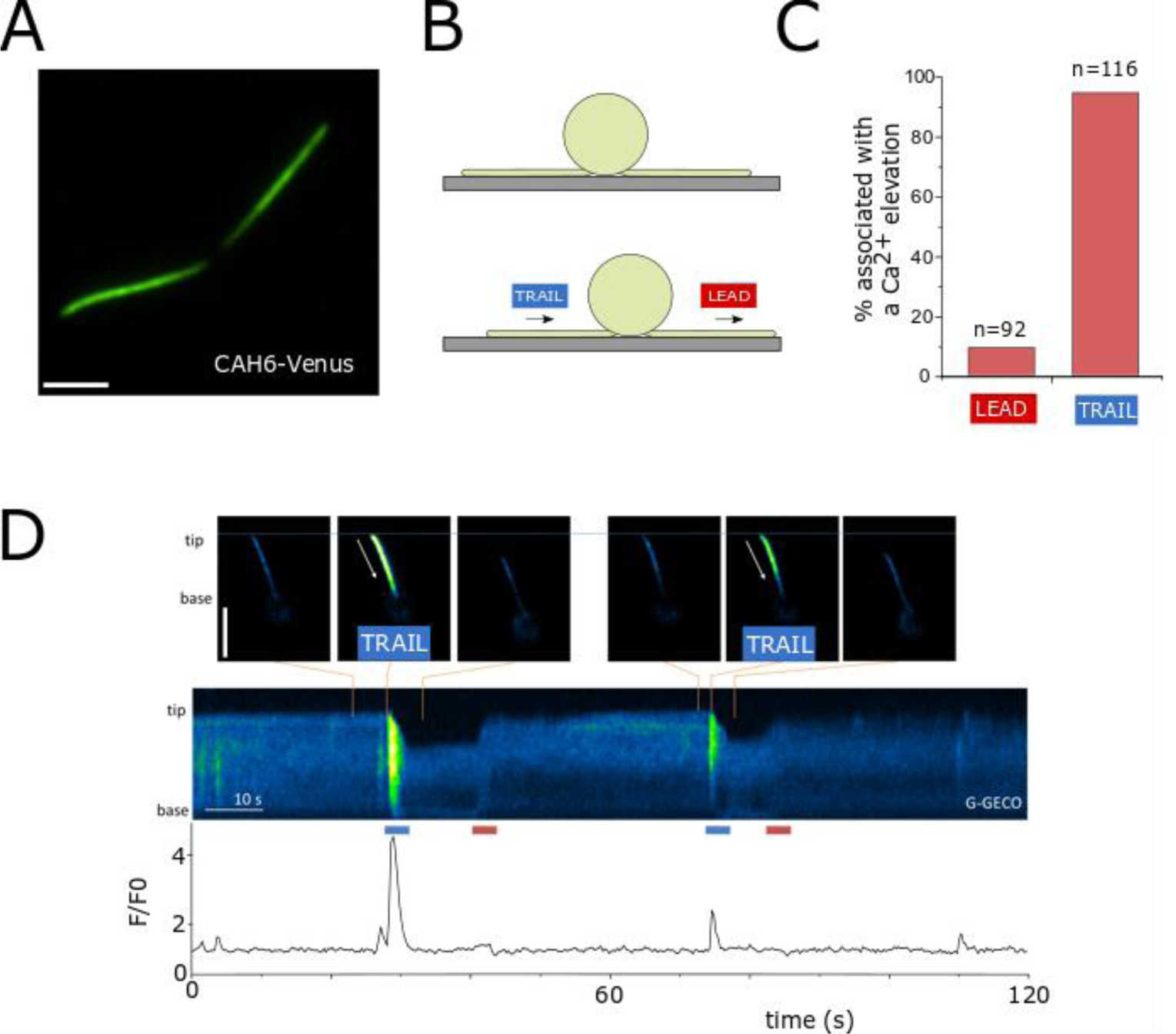
Development of a flagella-targeted calcium sensor in *Chlamydomonas*. **A)** Adherent flagella in a cell expressing CAH6-Venus, viewed by TIRF microscopy. Bar = 5 μm. **B**) *Chlamydomonas* cells with adherent flagella. Gliding movement is powered by retrograde IFT motors in the leading flagellum, dragging the trailing flagellum. Bar = 5 μm **C**) Quantitation of [Ca^2+^]_fla_ elevations associated with flagella movements. The percentage of gliding movements (leading and trailing) that coincide with a [Ca^2+^]_fla_ elevation. **D**) Kymograph illustrating flagella Ca^2+^ ([Ca^2+^]_fla_) elevations during gliding motility in wild type strain expressing CAH6-G-GECO (GG-WT strain). Significant [Ca^2+^]_fla_ elevations are associated with dragging movements in trailing flagella but not with forward movement in leading flagella.

We next examined the influence of different surface properties on gliding motility and flagella Ca^2+^ signalling. On non-treated glass coverslips, [Ca^2+^]_fla_ elevations in adherent flagella were associated primarily with flagella movement (Fig3A). Treating the glass coverslip with 0.1% poly-L-lysine greatly reduced gliding motility and flagella lifting, suggesting that the flagella are unable to overcome the increased adhesion to the surface. The immobilised adherent flagella exhibited highly repetitive [Ca^2+^]_fla_ elevations, with frequencies up to 0.57 Hz (Fig 3B). 63.6% of flagella on poly-lysine treated coverslips exhibited Ca^2+^ elevations at a frequency greater than 0.17 Hz, compared to 8.3% of flagella on untreated coverslips (Fig 3C). The repetitive [Ca^2+^]_fla_ elevations were not associated with motility, but occurred in immobilised flagella that were strongly adhered to the poly-lysine treated surface. Removing external Ca^2+^ strongly inhibited the repetitive [Ca^2+^]_fla_ elevations, indicating that they require Ca^2+^ influx across the flagella membrane (Fig 3D).

**Figure 3:**
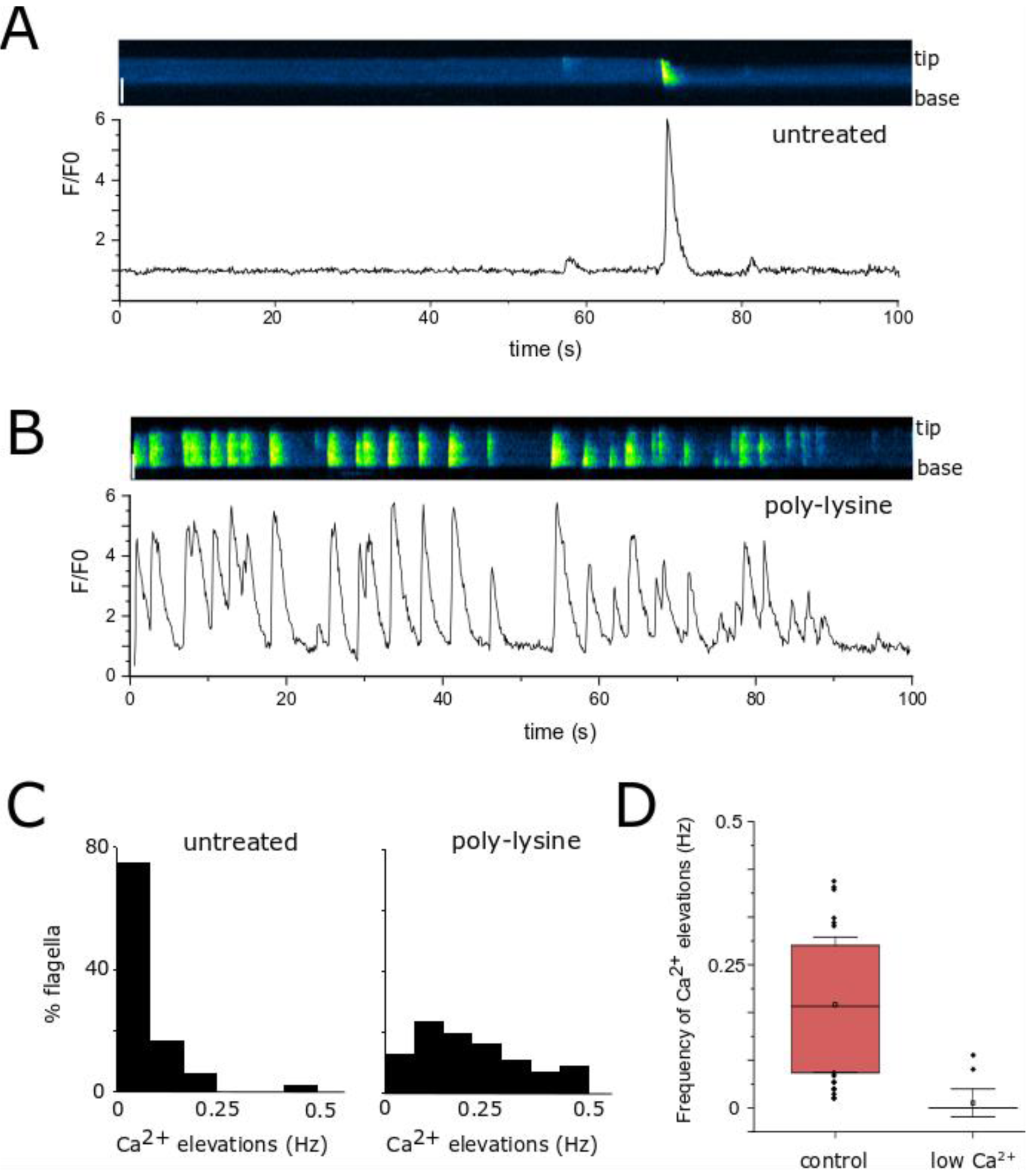
Increased adhesion results in high frequency Ca^2+^ spiking within flagella. **A)** [Ca^2+^]_fla_ elevations in a flagellum adhering to a non-treated glass coverslip in GG-IFT54MS strain expressing CAH6-G-GECO and IFT54-mScarlet. Bar = 5 μm. **B)** [Ca^2+^]_fla_ elevations in a flagellum adhering to a coverslip treated with 0.1% poly-lysine. Bar = 5 μm. **C)** Frequency histograms showing the percentage of flagella exhibiting different frequencies of [Ca^2+^]_fla_ elevations. Bin size = 0.083Hz, (n = 48 flagella on untreated surface, 55 flagella on poly-lysine). **D**) Repetitive [Ca^2+^]_fla_ elevations require external Ca^2+^. Boxplots showing the frequency of [Ca^2+^]_fla_ elevations in GG-IFT54MS cells after 20 minutes in different concentrations of external Ca^2+^ (free Ca^2+^ calculated as 300 μM or <1 μM). n = 34 and 21 flagella respectively.

### [Ca^2+^]_fla_ elevations coincide with the clearance of paused distal IFT accumulations

Expression of G-GECO in the IFT54-mScarlet reporter strain in combination with the immobilisation of flagella on poly-lysine treated coverslips enabled us to perform a detailed study of the interaction between [Ca^2+^]_fla_ elevations and distinct IFT processes. Flagella exhibiting a very high frequency of [Ca^2+^]_fla_ elevations (>0.25 Hz) were excluded from this initial analysis to prevent false correlations with high frequency IFT events (e.g. entry of IFT trains occurs at 1-1.3 Hz) ^32^. The entry of anterograde IFT trains into the flagellum, the arrival of anterograde IFT trains at the flagella tip and the departure of retrograde IFT trains from the flagella tip did not show a close relationship with [Ca^2+^]_fla_, as these IFT events occurred both in the presence and absence of [Ca^2+^]_fla_ elevations (Fig 4A-B). We also found no evidence for a correlation between elevated [Ca^2+^]_fla_ and the pausing or restarting of anterograde IFT trains. We were unable to clearly resolve the timing at which individual retrograde IFT trains paused, although it was clear that the characteristic distal IFT accumulations occur in periods lacking [Ca^2+^]_fla_ elevations (Fig 4A). However, in contrast to all other IFT events noted above, the restart of paused retrograde IFT trains showed a close association with elevated [Ca^2+^]_fla_ (Fig 4A-B). 90.2% of these events coincided with the onset of a [Ca^2+^]_fla_ elevation (defined as occurring within 0.5 s, n=41 events) (Fig 4C-D). Other IFT events only rarely coincided with the onset of [Ca^2+^]_fla_ elevations.

**Figure 4:**
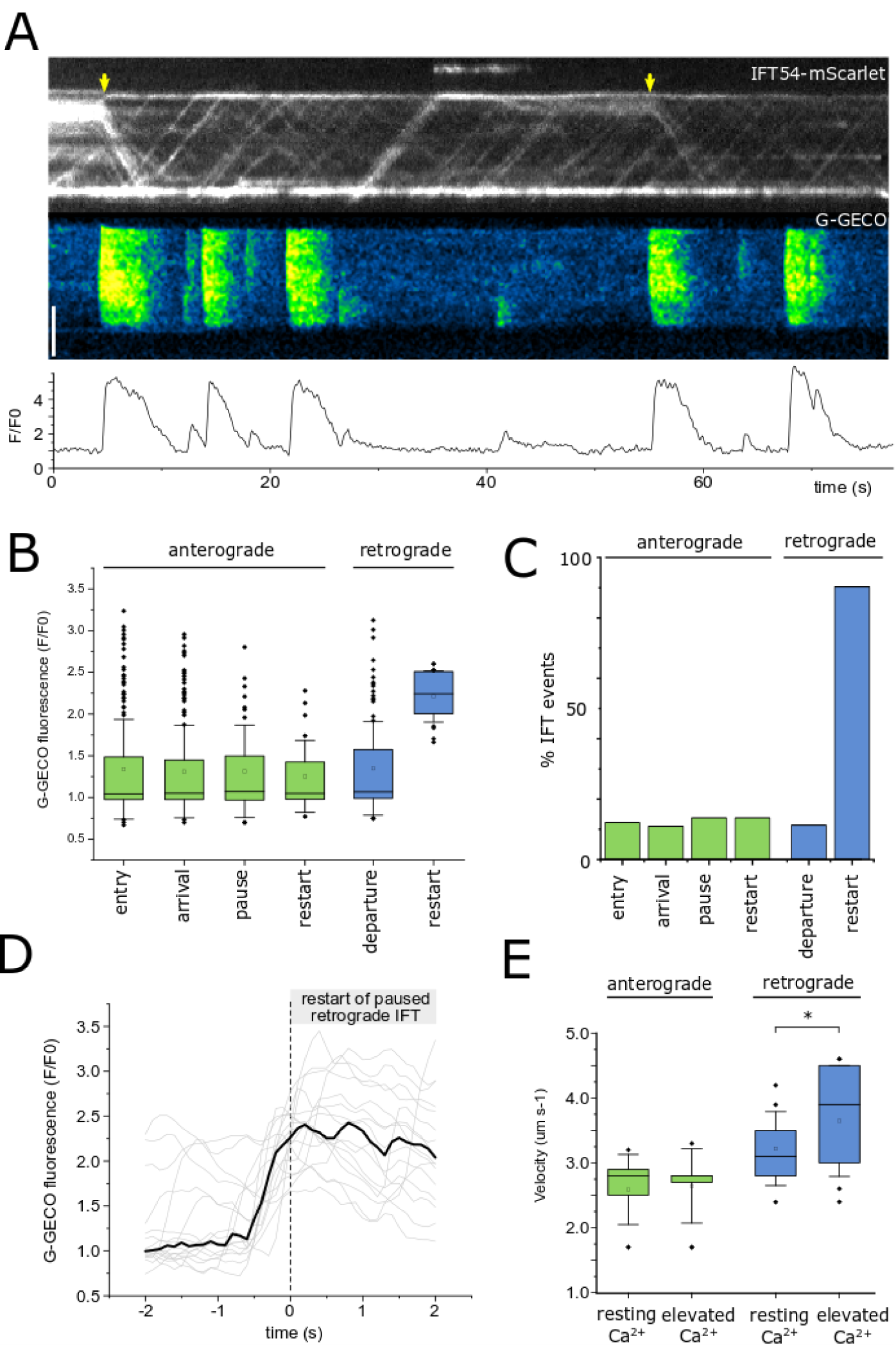
[Ca^2+^]_fla_ elevations coincide with retrograde movement of distal IFT accumulations. **A)** Simultaneous imaging of G-GECO and IFT54-mScarlet. The kymographs indicate the movement of IFT54 (upper) and changes in [Ca^2+^]_fla_ (lower) in a flagellum adhering to poly-lysine treated coverslip. Yellow arrow indicate movement of paused retrograde IFT trains. Bar = 5 μm. B) Box plot showing [Ca^2+^]_fla_ (displayed as relative G-GECO fluorescence) at the time of each IFT event. Six categories of IFT events were identified that could be clearly measured by TIRF microscopy. For anterograde IFT trains these were: entry into the flagellum, arrival at the flagella tip, pausing and restarting of paused anterograde IFT. For retrograde IFT, we identified trains departing from the flagella tip and the restarting of paused retrograde IFT. n = 154, 133, 32, 22, 92 and 17 IFT events for each category respectively. Box = median and 25-75% confidence interval, mean is open square, error bars represent standard deviation. C) Analysis of anterograde and retrograde IFT events that coincide with [Ca^2+^]_fla_ elevations. The frequency histogram illustrates the percentage of each IFT event that was observed to coincide with the initiation of a [Ca^2+^]_fla_ elevation. n = 1023, 82, 29, 29, 773, 41 IFT events for each category respectively. **D**) Timing of [Ca^2+^]_fla_ elevations associated with restart of paused retrograde IFT. 17 events are shown from 6 individual cells. Bold line = median trace, individual traces are shown in grey. **E**) Box plot showing the velocity of IFT trains during [Ca^2+^]_fla_ elevations (i.e. whilst [Ca^2+^]_fla_ was elevated above resting), relative to the velocity in the absence of a [Ca^2+^]_fla_ elevation. n = 9 flagella, with 200-410 IFT trains analysed for each category. Box = median and 25-75% confidence interval, mean is open square, error bars represent standard deviation. * denotes *p*<0.05 paired t-test.

In flagella exhibiting periodic [Ca^2+^]_fla_ elevations (0.1-0.25 Hz), the mean velocity of anterograde IFT trains was not influenced by elevated [Ca^2+^]_fla_, but there was a slight increase (+12.4 %) in the velocity of retrograde IFT trains in the presence of elevated [Ca^2+^]_fla_ (Fig 4E). This may indicate a reduction in transient pausing of retrograde IFT trains during [Ca^2+^]_fla_ elevations.

We conclude that the primary role of [Ca^2+^]_fla_ elevations in adherent flagella is to initiate the movement of paused retrograde IFT trains. We find no evidence for a direct requirement for [Ca^2+^]_fla_ elevations in the entry and turnaround of IFT trains, or in the pausing of anterograde IFT trains, suggesting that these processes are not directly regulated by the observed [Ca^2+^]_fla_ elevations, although our analysis does not rule out a role for Ca^2+^ in these processes via alternative mechanisms. For example, Ca^2+^-dependent protein kinases have been implicated in the dissociation of kinesin from IFT particles at the flagella tip during IFT turnover ^13^. As gradients in resting cytosolic Ca^2+^ can be observed in some cells, notably in polarised plant cells such as root hairs or pollen tubes ^33, 34^, we investigated whether resting [Ca^2+^]_fla_ differed at the flagella tip. The median intensity of CAH6-G-GECO or CAH6-Venus (as a non-Ca^2+^-responsive control) did not differ along the length of the flagellum (Supplementary Fig 2). We also examined cells that had been biolistically loaded with the Ca^2+^-responsive dye Oregon Green BAPTA dextran (OGB) and the non-Ca^2+^-responsive dye Texas Red dextran (TR), enabling ratiometric imaging of [Ca^2+^]_fla_. The ratio of OGB/TR was similar along the length of the flagellum, indicating that resting [Ca^2+^]_fla_ is not consistently elevated at the flagella tip relative to the rest of the flagellum.

### Repetitive [Ca^2+^]_fla_ elevations prevent distal IFT accumulations in adherent flagella

Comparison of flagella on untreated and poly-lysine treated surfaces indicated that the nature of IFT differed markedly. On a poly-lysine treated surface, where flagella exhibit repetitive Ca^2+^ elevations, the movement of IFT trains was more regular, with a higher frequency of anterograde and retrograde IFT trains (Fig 5A-C). Notably, during periods of high frequency [Ca^2+^]_fla_ elevations, significant distal accumulations of retrograde IFT trains were not observed, whereas anterograde IFT trains continued to pause and restart. These observations suggest that repetitive [Ca^2+^]_fla_ elevations act to prevent the distal accumulations of paused retrograde IFT trains. Examination of flagella exhibiting intermittent periods of repetitive Ca^2+^ fla elevations revealed that the transition between these patterns of IFT could be very rapid (Supplementary Fig 3; Video 2). Experimentally inhibiting Ca^2+^ signalling using a perfusion system to remove external Ca^2+^ resulted in the rapid cessation of [Ca^2+^]_fla_ elevations and the accumulation of IFT trains in the distal region (Supplementary fig 4). This coincided with a clear reduction in the frequency of retrograde IFT trains.

**Figure 5:**
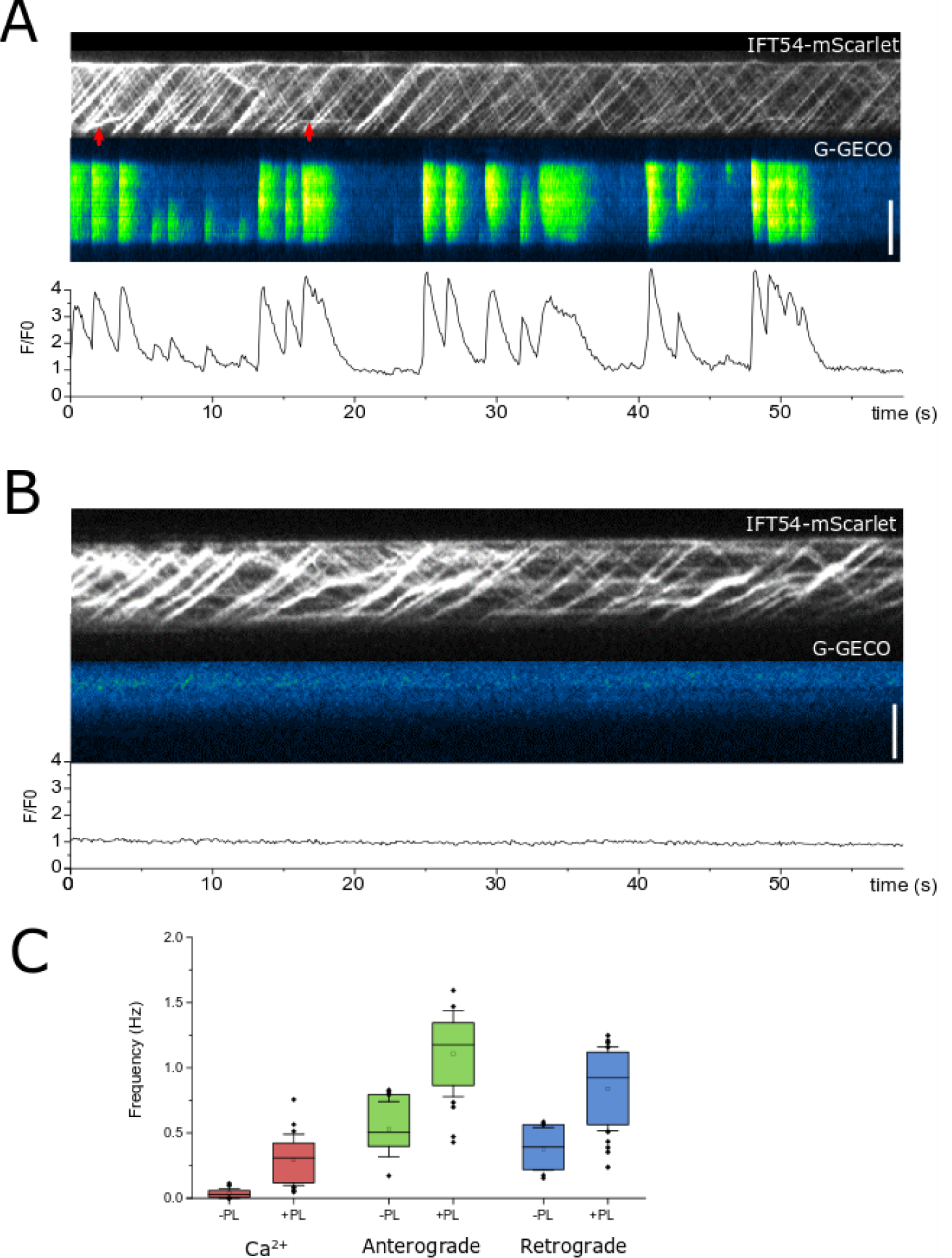
Flagella exhibiting high frequency [Ca^2+^]_fla_ elevations do not exhibit IFT accumulations. **A**) Simultaneous TIRF imaging of [Ca^2+^]_fla_ and IFT54-mScarlet on a poly-lysine treated surface. Flagella exhibit highly repetitive [Ca^2+^]_fla_ elevations, with a high frequency of anterograde and retrograde IFT. Pausing of anterograde IFT trains can still be observed (red arrows). **B**) As in (A) but on an untreated surface. **C**) Box plots showing the frequency of [Ca^2+^]_fla_ elevations, anterograde IFT and retrograde IFT on untreated and poly-lysine treated (+PL) surfaces (n=11 and 20 flagella respectively). Box = median and 25-75% confidence interval, mean is open square, error bars represent standard deviation.

We have previously shown that removing external Ca^2+^ for longer periods (>5 minutes) results in the substantial overaccumulation of IFT20-mCherry in adherent flagella ^20^ and confirmed that this was also the case for IFT54-mScarlet and D1bLIC-GFP (Supplementary Fig 4A-B). Retrograde IFT is largely absent from these flagella, although anterograde IFT can still be observed, albeit with a reduced frequency and velocity. As [Ca^2+^]_fla_ elevations have no short term impact on the pausing of anterograde IFT trains or their velocity (Fig 4), the effect of removing Ca^2+^ on anterograde IFT likely results from the overaccumulation of retrograde IFT trains. Anterograde IFT trains frequently paused or slowed in the vicinity of large accumulations of IFT trains, suggesting that they are impeded by the paused retrograde IFT trains. This conclusion is seemingly at odds with the finding that anterograde and retrograde IFT trains travel on distinct microtubules and do not collide ^27^. We therefore examined the nature of these distal accumulations by transmission electron microscopy (TEM) of adherent flagella. After the removal of external Ca^2+^ for 5 minutes, TEM of adherent flagella revealed large accumulations of IFT particles in a prominent swelling near the distal tip (Supplementary Fig 4C). The accumulated IFT trains represent a similar phenotype to the flagella of strains defective in retrograde IFT, such as mutants in cytoplasmic dynein ^30, 35^. We propose that prolonged inhibition of Ca^2+^ signalling results in retrograde IFT trains accumulating behind the retrograde IFT trains that have already paused. In the absence of a mechanism to clear the paused retrograde IFT trains, they can accumulate to an extent where they significantly impede the progress of anterograde IFT trains, even if these IFT trains move along a distinct microtubule track.

### Detachment from a surface requires external Ca^2+^

Our results show that [Ca^2+^]_fla_ elevations are required to regulate the accumulation of paused retrograde IFT trains. This process is essential for gliding motility ^20^, but may also modulate flagella adhesion to a surface. Flagella lifting is often initiated by axonemal bending in the non-adherent region of the flagellum, resulting in the flagellum being dragged towards the cell body so that only the flagella tips remain adherent. We found that flagella lifting was strongly associated with [Ca^2+^]_fla_ elevations, with both the GG-WT and GG-IFT54-MS strains routinely exhibiting a [Ca^2+^]_fla_ elevation with each lifting event (Supplementary Fig 5A-B). Detailed examination of the timing of lifting events revealed that in all cases the [Ca^2+^]_fla_ elevation immediately preceded the initial movement of the flagellum towards the cell body (Supplementary Fig 5C). [Ca^2+^]_fla_ elevations associated with lifting and gliding are therefore both linked to the dragging of flagella along a surface, suggesting that they are caused by an increase in membrane tension. We propose that the [Ca^2+^]_fla_ elevations act to release FMG-1B from paused retrograde IFT trains, which removes resistance to the pulling force generated by bending the axoneme and allows the flagella to be withdrawn towards the cell body and lifted from the surface.

We next examined whether Ca^2+^-dependent removal of paused retrograde IFT trains was also required for light-modulated flagella detachment. *Chlamydomonas* cells (strain SAG-11-32b) suspended on a micropipette detach their flagella from a surface within 30 s after the removing blue light ^22^. We found that removing blue light from adherent IFT54-MS cells also promoted flagella detachment after 30 seconds, with flagella either remaining only attached at the flagella tips or lifting from the surface entirely (11 out of 14 cells) (Supplementary Fig 5D). Flagella movements prior to lifting were preceded by the removal of paused retrograde IFT trains (Supplementary Fig 5E). As G-GECO requires blue light excitation, it was not possible to measure [Ca^2+^]_fla_ directly during these movements. However, removing blue light in the absence of external Ca^2+^ did not result in flagella lifting or the removal of paused retrograde IFT trains (0 out of 17 cells) (Supplementary Fig 5F). Note that non-adherent cells were able to swim in the absence of Ca^2+^, indicating that inhibition of Ca^2+^ signalling did not act to simply immobilise flagella. The results indicate that light-dependent flagella detachment requires the Ca^2+^-dependent release of FMG-1B from paused retrograde IFT trains. This allows FMG-1B to move in an unrestricted manner in the flagella membrane, facilitating the lifting movements.

### [Ca^2+^]_fla_ elevations have distinct spatial properties

The [Ca^2+^]_fla_ elevations observed in *Chlamydomonas* not only show complex temporal patterns, but also distinct spatial properties. The majority of [Ca^2+^]_fla_ elevations spanned the entire length of the flagellum. However, in cells exhibiting lower frequency Ca^2+^ elevations (<0.25 Hz), 35.7% of the [Ca^2+^]_fla_ elevations were restricted to either the proximal or the distal region of the flagellum (Fig 6A-B). In addition, many of the [Ca^2+^]_fla_ elevations that did span the length of the flagellum did not rise simultaneously along the length of the flagellum. 28.1% of [Ca^2+^]_fla_ elevations exhibited a difference >0.2 s in onset between the distal and proximal regions (Fig 6C). Closer examination of these signalling events indicated that they resemble a Ca^2+^ wave that propagates along the length of the flagellum in either direction, with 43.8% propagating in a retrograde direction and 56.2% propagating in an anterograde direction. The mean rate of propagation was 28.7 ± 4.3 μm s^-1^ (n=10, ± s.e.) (Fig 6D).

**Figure 6:**
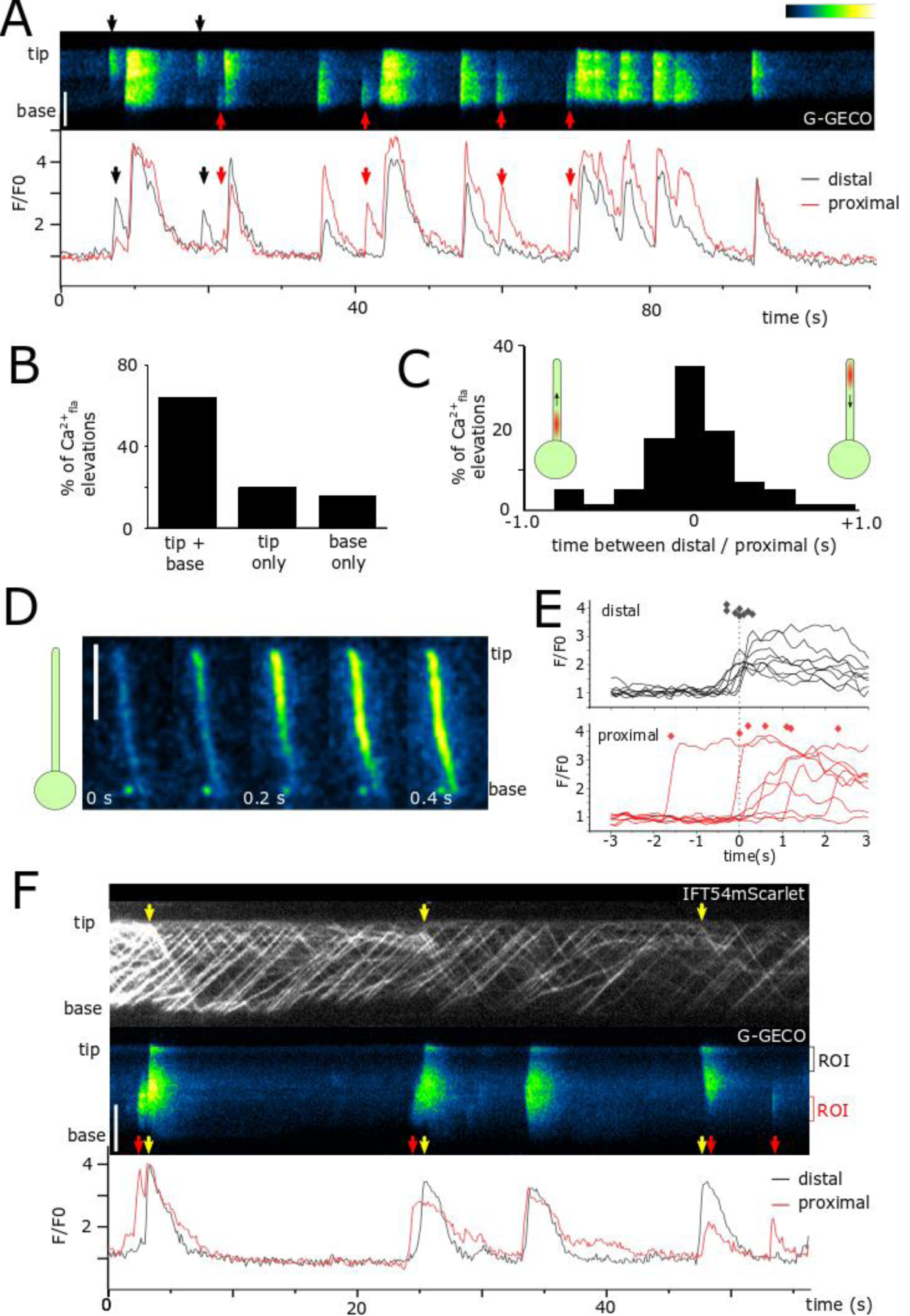
[Ca^2+^]_fla_ elevations have distinct spatial properties. **A**) Kymograph showing the presence of localised Ca^2+^ elevations in GG-IFT54MS strain restricted to either the distal region (black arrow) or proximal region (red arrow). Bar = 5 μm. **B**) Frequency histogram displaying the percentage of [Ca^2+^]_fla_ elevations that occur across the whole flagellum or are restricted to either the distal or proximal regions. n = 250 [Ca^2+^]_fla_ elevations from 7 flagella. **C**) Frequency histogram displaying the time difference between the rise time of [Ca^2+^]_fla_ elevations in the distal region compared to proximal region for Ca^2+^ elevations that span the whole flagellum. Positive numbers reflect [Ca^2+^]_fla_ elevations that rise first in the distal region and spread to the proximal region, whereas negative numbers indicate [Ca^2+^]_fla_ elevations that initially arise in proximal regions. n = 57 elevations from 9 flagella. Bin size = 0.2 seconds. **D**) Image series displaying a [Ca^2+^]_fla_ elevation that propagates from the distal region. Bar = 5 μm. **E**) Timing of [Ca^2+^]_fla_ elevations in the distal and proximal regions of the flagellum, displayed relative to the onset of movement of paused retrograde IFT trains in the distal region (dotted line). Diamonds represent the onset of each [Ca^2+^]_fla_ elevation defined as the greatest rate of change in fluorescence. Whilst [Ca^2+^]_fla_ elevations in the distal region correspond closely to the movement of IFT trains, [Ca^2+^]_fla_ elevations in the proximal region exhibit variable timing or are absent. **F**) Simultaneous imaging of [Ca^2+^]_fla_ and IFT54-mScarlet indicates that movement of paused retrograde IFT trains in the distal region of the flagellum (yellow arrow) coincides with co-localising [Ca^2+^]_fla_ elevations. Red arrows show [Ca^2+^]_fla_ elevations restricted to the proximal region. Bar = 5 μm.

We next examined how propagating [Ca^2+^]_fla_ elevations influenced the movement of paused IFT trains to determine whether Ca^2+^ acted in a local or global manner within the flagellum. Irrespective of whether the [Ca^2+^]_fla_ wave initiated in the distal or proximal region, we found that the movement of paused retrograde IFT trains in the distal region corresponded closely to the time at which [Ca^2+^]_fla_ became elevated in the distal region, rather than the proximal region (Fig 6E-F). This indicates that [Ca^2+^]_fla_ elevations act locally, requiring co-localisation with the paused retrograde IFT trains in order to initiate their movement.

### [Ca^2+^]_fla_ elevations disrupt microsphere movement along the flagellum

Our results strongly support a role for Ca^2+^ in regulating the interaction of FMG-1B with IFT particles. This interaction can be readily visualised by following the movement of fluorescent microspheres along the flagellum ^17^. Through simultaneous imaging of fluorescent microspheres and IFT27-GFP, we observed that microspheres stopped moving in a particular direction when the microsphere switched to an IFT train moving in the opposite direction or reached the end of the flagellum (Fig 7A). However, on some occasions the microspheres could be observed to dissociate from the IFT train (anterograde or retrograde) and exhibit diffusional movement in the flagella membrane. We hypothesised that these events represented a disruption of the interaction between the IFT particle and FMG-1B, which could be mediated by [Ca^2+^]_fla_ elevations. We therefore next observed microsphere movements along a flagellum whilst simultaneously monitoring [Ca^2+^]_fla_. 12 out of 25 flagella exhibiting IFT-driven microsphere movement also exhibited a [Ca^2+^]_fla_ elevation during this period. In each of these flagella, the [Ca^2+^]_fla_ elevation coincided directly with the cessation of microsphere movement, with the microsphere either subsequently exhibiting diffusional movement or becoming detached from the flagellum entirely (Fig 7B-C; Video 3). We attribute the latter events to a strong attraction between the microsphere and the poly-lysine treated surface. Moreover, we did not observe microspheres moving on flagella that exhibited high frequency [Ca^2+^]_fla_ elevations. We conclude that [Ca^2+^]_fla_ elevations act directly to disrupt the interaction between IFT trains and FMG-1B.

**Figure 7:**
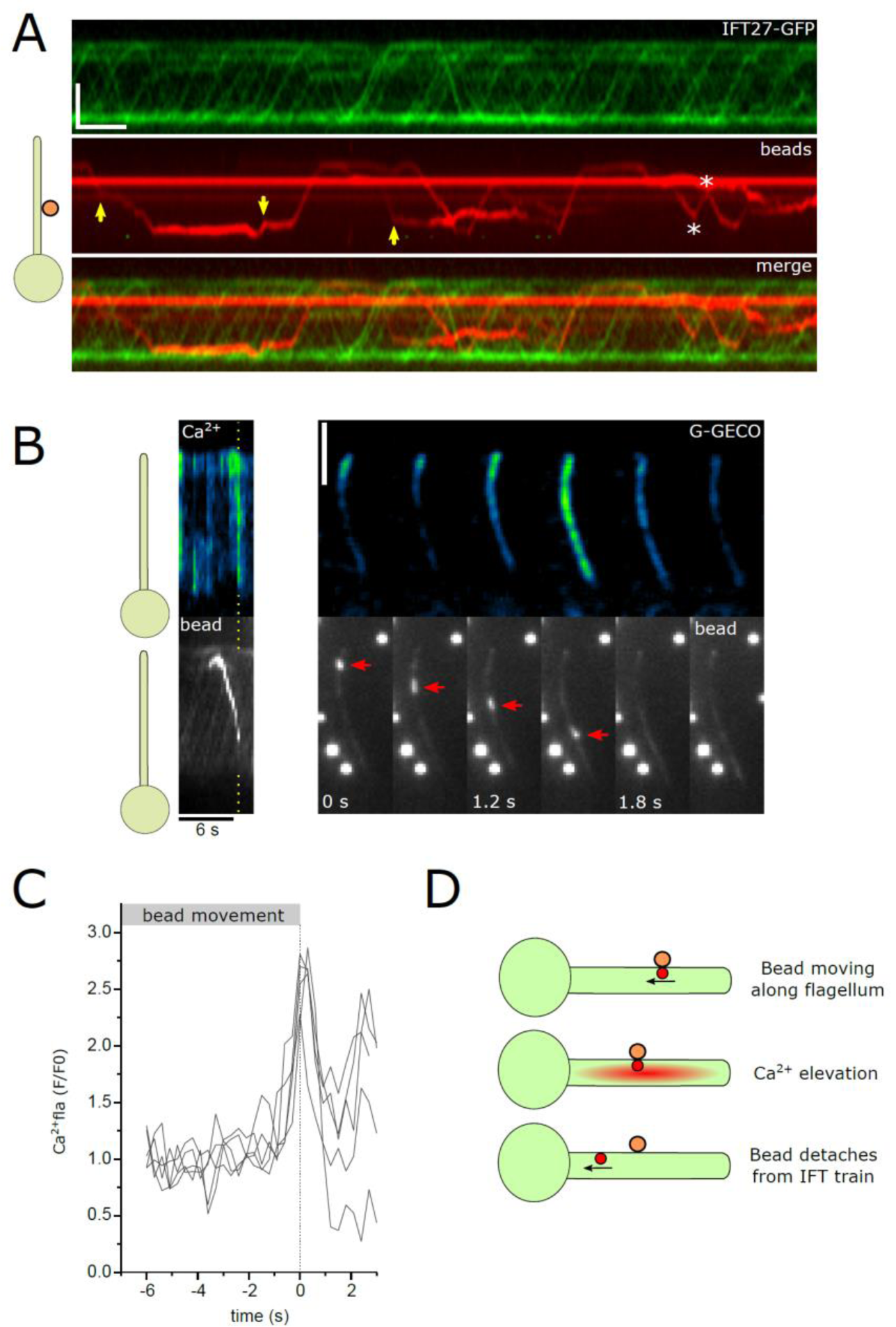
[Ca^2+^]_fla_ elevations inhibit movement of microspheres along the flagellum. **A**) Kymograph displaying simultaneous TIRF imaging of IFT27-GFP and fluorescent microspheres. Microspheres moving in both anterograde and retrograde directions co-localise with IFT trains. Yellow arrows indicate events where microspheres dissociate from the IFT train and exhibit diffusive movement. Asterisks indicate events where microspheres rapidly change direction by associating to IFT trains moving in the opposite direction. Bar = 5 μm and 10 s. **B**) Kymograph displaying simultaneous TIRF imaging of [Ca^2+^]_fla_ and fluorescent microspheres. The image sequence demonstrates that the [Ca^2+^]_fla_ elevation coincides directly with the end of movement of the microsphere (arrowed), which in this case is lost from the flagellum. Bar = 5 μm. **C**) [Ca^2+^]_fla_ elevations in flagella displaying IFT-driven microsphere movement. [Ca^2+^]_fla_ traces are shown relative to the cessation of IFT-driven microsphere movement (n=5 flagella). **D**) Schematic displaying the proposed signalling events involved in the detachment of microspheres from IFT trains.

## Discussion

The mechanisms regulating the interactions between IFT complexes and their cargo proteins underpin many aspects of ciliary function ^36^. Our results show that Ca^2+^ signalling plays an important role in regulating transient interactions between the flagella membrane glycoprotein FMG-1B and the IFT complex in adherent *Chlamydomonas* flagella. The primary role of the [Ca^2+^]_fla_ elevations is to disrupt the interaction between FMG-1B and paused retrograde IFT trains, which allows the retrograde IFT trains to return to the cell body and releases FMG-1B to move freely in the flagella membrane. We did not find evidence for a requirement for [Ca^2+^]_fla_ elevations in regulating other aspects of IFT, for example by promoting the pausing of anterograde or retrograde IFT trains or directly influencing their movement. Chien et al ^2^ found that the removal of external Ca^2+^ increased the residency of IFT trains at the flagella tip, suggesting that Ca^2+^ may have a regulatory role in the assembly or departure of retrograde trains. We were not able to directly assess the turnaround of IFT trains in our analyses. However, we did observe that retrograde IFT trains continue to accumulate in the absence of external Ca^2+^, indicating that IFT turnaround can continue following the inhibition of [Ca^2+^]_fla_ elevations. Moreover, we observed that many retrograde IFT trains paused directly in the vicinity of the flagella tip, which may influence measurement of turnover.

Whilst the interaction between the IFT particle and FMG-1B is implied from the coordinated movement of fluorescent microspheres and IFT trains ^17^, biochemical characterisation of this interaction is lacking. Recent characterisation of an *fmg-1b* mutant line ^37^ supports previous observations that FMG-1B is the primary protein in the flagellum mediating contact with solid substrates ^38, 39^, although an additional flagella membrane glycoprotein, FAP113, was recently identified through its interactions with microspheres ^40^. It seems likely that surface contact, which induces cross-linking of FMG-1B ^41^, acts to promote an interaction between FMG-1B and retrograde IFT trains, as retrograde IFT trains do not accumulate in non-adherent flagella ^20^. We propose that elevations in [Ca^2+^]_fla_ in the vicinity of the paused IFT train likely lead to a direct conformational shift in a Ca^2+^-sensitive protein within the putative IFT/FMG-1B complex, disrupting the interaction. This mechanism would be analogous to the regulated movement of mitochondria along microtubules in neurons and other animal cell types, where the interaction between the mitochondrion and kinesin or dynein is mediated by the Ca^2+^-binding Rho-GTPase Miro ^42^. Elevated cytosolic Ca^2+^ triggers a conformational change in Miro, which possesses two Ca^2+^-binding EF-hands, leading to the dissociation of the mitochondrion from the microtubule motor complex ^43^.

[Ca^2+^]_fla_ elevations in adherent *Chlamydomonas* flagella were primarily observed in flagella under tension, i.e. immediately prior to trailing or lifting movements, and flagella that were not moving usually did not exhibit [Ca^2+^]_fla_ elevations. In contrast, flagella immobilised on a highly adherent surface often exhibited high frequency repetitive [Ca^2+^]_fla_ elevations. This suggests that immobilisation in this manner also causes membrane tension, triggering the repetitive [Ca^2+^]_fla_ elevations. Our results indicate that the primary role of [Ca^2+^]_fla_ elevations in adherent flagella relates to modulating their adhesion to the surface. When not in contact with anterograde or retrograde IFT trains, FMG-1B can diffuse freely along the flagellum, as demonstrated by microsphere movements ^37^. This allows an adherent flagellum to slide along a surface whilst maintaining contact with it. In a trailing flagellum, some of the FMG-1B is shed onto the surface, leaving a ‘snail trail’ of adhesive glycoprotein, which allows gliding motility to continue beyond a single flagellum length ^40^ (Supplementary Fig 6). However, when FMG-1B becomes bound to retrograde IFT trains, the dynein motors act to pull the flagellum forward. The flagellum is therefore resistant to any movement in the direction of the cell body, unless the interactions between FMG-1B and the IFT trains are disrupted. The resistance to the pulling force likely contributes to an increase in membrane tension that triggers [Ca^2+^]_fla_ elevations. The [Ca^2+^]_fla_ elevations therefore do not act to modify adhesion directly, but to allow unrestricted movement of FMG-1B along the membrane, which in turn allows the flagellum to be dragged along a surface to enable gliding motility or flagella lifting (Supplementary Fig 7). If the flagellum is immobilised on a highly adherent surface, the membrane tension cannot be relieved, leading to the highly repetitive [Ca^2+^]_fla_ elevations observed on poly-lysine treated surfaces. Although the repetitive [Ca^2+^]_fla_ elevations act to prevent the accumulation of paused retrograde IFT trains, in this situation the flagella remain immobilised due to the highly adherent nature of the surface itself.

Like many cilia, *Chlamydomonas* flagella are mechanosensitive and contain a number of candidate ion channels that may contribute to signalling in response to changes in membrane tension ^24, 44, 45^. RNAi knockdown of the transient receptor potential (TRP) channel TRP11 resulted in a defect in the mechanoshock response of swimming *Chlamydomonas* cells ^46^. *Chlamydomonas* flagella also possess a homologue of PKD2, which has been implicated in mechanosensation in mammalian cilia, although direct evidence for mechanosensory calcium signalling in primary cilia is lacking ^47, 48^. *Chlamydomonas* PKD2 localises to the flagella membrane, but is also bound to the axoneme and to extracellular glycoprotein polymers known as mastigonemes that resemble hair-like appendages along the distal regions of the flagella ^49, 50^. The tethering of PKD2 to mastigonemes and the axoneme supports a mechanosensory role in *Chlamydomonas* ^50^.

The development of flagella-localised Ca^2+^-reporters in *Chlamydomonas* should now aid the molecular identification of specific ion channels responsible for flagellar Ca^2+^ signalling in adherent flagella. However, the nature of flagella adhesion and membrane tension will be a critical consideration for future studies of flagella Ca^2+^ signalling, particularly when comparing phenotypes between strains. Gliding motility is often markedly different between strains and rapid modulation of flagella adhesion can occur within seconds even within individual cells ^22^. The *pkd2* mutants’ lack of mastigonemes, indicates that ion channel mutants may influence the arrangement of flagella glycoproteins ^50^. Whilst mastigonemes are not thought to be involved in microsphere movements or gliding motility ^51, 52^, it is clear that knockout of an ion channel could influence the processes that trigger [Ca^2+^]_fla_ elevations (i.e. flagella adhesion). Without direct measurements of the degree of adhesion and/or membrane tension in these mutant strains, it may be difficult to ascertain whether any effects on Ca^2+^ signalling result from direct mechanistic inhibition of [Ca^2+^]_fla_ elevations or relate to indirect effects on flagella adhesion.

Repetitive Ca^2+^ elevations can be observed in many excitable cell types in the presence of a continuous stimulus. In adherent *Chlamydomonas* flagella, the initial activation of a mechanosensitive ion channel may act to depolarise the flagella membrane, leading to activation of voltage-gated Ca^2+^ channels and a resultant action potential. If the stimulus persists (i.e. membrane tension), the voltage-gated Ca^2+^ channels can activate again following a refractory period, leading to a series of repetitive Ca^2+^ elevations. The presence of voltage-gated Ca^2+^ channels in *Chlamydomonas* flagella is well documented ^53, 54^ and we have previously observed transient depolarisations in the flagella membrane during gliding motility ^20^. The presence of localised or propagating [Ca^2+^]_fla_ elevations suggests that the mechanisms underlying flagella Ca^2+^ signalling may exhibit greater complexity. It is possible that the propagating [Ca^2+^]_fla_ elevations represent simple diffusion of Ca^2+^ in the flagella matrix following a localised [Ca^2+^]_fla_ elevation. In mammalian ependymal cells, Ca^2+^ elevations in the cytosol can diffuse along cilia at a rate of approximately 20 µm s^-1 55^. However, our previous studies suggest that Ca^2+^ elevations in the *Chlamydomonas* cytosol are not directly coupled to [Ca^2+^]_fla_ elevations ^20, 56^, which may indicate a greater degree of Ca^2+^ buffering in *Chlamydomonas* flagella. In addition, ion channels that are typically associated with the active propagation of cytosolic Ca^2+^ elevations in animal cells are found in *Chlamydomonas* flagella. In mammalian cells, inositol triphosphate receptors (IP_3_R) typically localise to endomembranes, where they contribute to cytosolic Ca^2+^ waves through Ca^2+^-activated Ca^2+^ release. A homologue of mammalian IP_3_Rs is present in *Chlamydomonas* flagella ^57^, which could potentially mediate propagation of localised [Ca^2+^]_fla_ elevations through Ca^2+^-activated Ca^2+^ influx.

It is becoming clear that the nature of ciliary Ca^2+^ signalling differs markedly between organisms and cell types ^58^. Rapid changes in flagella Ca^2+^ have a well characterised role in the swimming motility of *Chlamydomonas*, in addition to their role in gliding motility ^59^. Similar Ca^2+^-dependent signalling mechanisms regulate swimming of ciliates such as *Paramecium* ^60^ and repetitive [Ca^2+^]_fla_ elevations associated with chemotactic motile responses in metazoan sperm have also been studied extensively ^61^. Whilst it likely that flagella-localised voltage-gated Ca^2+^ channels contribute directly to [Ca^2+^]_fla_ elevations in *Chlamydomonas* and *Paramecium* ^54, 60^, their role in mammalian sperm flagella are less clear. Patch-clamp approaches in mammalian sperm suggest that the weakly-voltage-gated CatSper channels are primary contributors to flagella Ca^2+^ elevations, rather than voltage-gated Ca^2+^ channels (Cav) ^62^. Although CatSpers are also found in some protist lineages (e.g. glaucophytes), they are absent from *Chlamydomonas* ^63^. Furthermore, whilst voltage-gated Ca^2+^ channels are present in motile ependymal cilia of mammalian cells, ciliary Ca^2+^ in these cells appeared to be passively coupled to [Ca^2+^]_cyt_ rather than independently controlled ^55^. The nature of Ca^2+^ signalling in the non-motile primary cilia of mammalian cells is also highly distinct from the excitable cilia of unicellular protists, with primary cilia exhibiting a high resting Ca^2+^ determined primarily by the activity of the TRP channel PKD2L1 ^12^. Therefore, although Ca^2+^-dependent signalling mechanisms appear to be ubiquitous in cilia and flagella, the nature of these mechanisms and their underlying molecular components differ considerably between eukaryotes.

In summary, our results indicate that interactions between IFT trains and flagella membrane proteins can be directly influenced by rapid transient [Ca^2+^]_fla_ elevations. As FMG-1B is an abundant protein in *Chlamydomonas* flagella, these interactions can be readily observed through the movement of microspheres or gliding motility. However, it is likely that regulated transient interactions with IFT complexes also direct the movement of many other flagella membrane proteins. Future biochemical characterisation of the FMG-1B/IFT complex, and in particular the identification of the Ca^2+^-sensitive components that mediate the response to [Ca^2+^]_fla_ elevations, will help identify whether these mechanisms are conserved in other ciliated organisms.

## Materials and Methods

### Strains and culture conditions

*Chlamydomonas reinhardtii* cells were grown in standard Tris-acetate-phosphate (TAP) media at 21°C on a 16:8 hour light:dark cycle with a photosynthetic photon flux density of 80 μmol m^-2^ s^-1^. Wild type strain CC-5325, D1bLIC-GFP (CC-4488) ^28^ and KAP-GFP (CC-4296 - *fla3-1* background ^29^) were obtained from the Chlamydomonas Resource Centre (http://chlamycollection.org/). *Chlamydomonas* strains expressing CAH6-Venus and IFT27-GFP (CC125 background) were obtained from Luke Mackinder (York) and Joel Rosenbaum (Yale) respectively ^31, 64^. IFT20-mCherry (null *ift20* background ^65^) and the IFT54-mScarlet reporter strain (IFT54-MS) were provided by Karl Lechtreck (University of Georgia). The IFT54-MS strain was constructed by expressing IFT54-mScarlet in an *ift54-2* background ^32^. The IFT54-mScarlet vector was made by cutting out mNeonGreen from pBR25-mNG-IFT54 using *XhoI* and *BamHI* and replaced with a PCR product encoding mScarlet-I using the mammalian codon usage trimmed with the same enzymes ^32^.

### Development of a flagella-targeted calcium sensor

The flagella targeted calcium reporter was prepared by modifying the pLM005-CAH6-Venus vector used to express the flagella-localised carbonic anhydrase CAH6 ^31^. Codon-optimised G-GECO1.1 was synthesised (Genscript) and introduced into the pLM005-CAH6-Venus expression vector via the sites *BglII* and *EcoRI*, resulting in a C-terminal fusion of G-GECO to CAH6, separated by a short amino acid linker sequence. The GG-WT and GG-IFT54-MS strains were produced by transforming *C. reinhardtii* strains CC5325 (wild type) and IFT54-MS by electroporation using a NEPA21 Super Electroporator (NepaGene, Japan) ^66^. Cells were grown to a cell density of 1-2×10^7^ cells mL^-1^. The cultured cells were collected by centrifugation (2000 rpm, 3 min) and re-suspended in TAP media containing 40 mM sucrose to a final density of 1-1.5×10^8^ cells mL^-1^. 1×10^7^ cells and 350 ng of plasmid DNA (pLM005-CAH6-G-GECO construct linearized with *EcoRV*) were suspended in a total volume of 20-30 μL and placed into an electroporation cuvette (2 mm). The measured value of electric impedance was within 500-650 Ω. Electroporation utilised a first pulse (Pp) at 250 V with 8 ms pulse length, 50 ms pulse interval and 40% decay rates. The second pulse consisted of multiple transfer pulses (Tp) of 20 V with 50 ms pulse length, 50 ms pulse interval and 40% decay rate (2 iterations). Cells were was transferred into 10 mL TAP media containing 40 mM sucrose and incubated in dim light (2-3 µmol photons m^-2^ s^-1^) for 24 h. The cells were collected by centrifugation (2000 rpm, 3 min) and plated onto 1.5 % agar TAP plate containing 10 µg mL^-1^ hygromycin B (Roche) and 20 µg mL^-1^ of paromomycin sulfate (Fisher BioReagents).

### Biolistic loading of Ca^2+^ dyes

For experiments where flagella Ca^2+^ was measured using fluorescent dyes, dyes were loaded into cells using a biolistic approach ^20^. 0.9 mg of 0.6 µm diameter gold microcarriers (Bio-Rad) were coated in 40 µg Oregon green-BAPTA dextran (10000 MW) and 24 µg Texas red dextran (10000 MW) (Invitrogen). Oregon Green-BAPTA is Ca^2+^-responsive, whereas Texas Red acts as a non-responsive reference dye. Texas Red was omitted for simultaneous imaging of Ca^2+^ and IFT20-mCherry. *Chlamydomonas* cells were harvested in mid-exponential phase by gentle centrifugation (400 *g* for 3 min) and resuspended in loading buffer (10 mM HEPES pH 7.4, 20 µM K^+^ glutamate and 50 mM sorbitol). 5 x 10^6^ cells were spread onto a 0.45 µm nitrocellulose membrane filter (Millipore) and loaded biolistically using a PDS-1000 delivery system (Bio-Rad) (1100 psi rupture disc). After loading, cells were washed and resuspended in TAP media and allowed to recover for at least 2 hours prior to imaging.

### TIRF imaging of flagella

For all experiments, *Chlamydomonas* cells were suspended in a HEPES/NMG buffer consisting of 5 mM HEPES, 1 mM KCl, 1 mM HCl, 500 µM CaCl_2_, 200 µM EGTA (free Ca^2+^ 301 µM) and pH adjusted to 7.4 with N-methyl-D-glucamine [6]. To remove external Ca^2+^, cells were perfused with a Ca^2+^-free buffer in which CaCl_2_ had been omitted. Cells were added to 35 mm glass bottom dishes (In Vitro Scientific), which in some cases were treated with 0.1% poly-lysine to promote flagella adhesion. TIRF microscopy was performed at room-temperature using a Nikon Eclipse Ti with 100x (1.49 NA) TIRF oil immersion objective and an EM-CCD camera (Photometrics Evolve). TIRF was achieved via through-the-lens laser illumination. Wavelength compensation was not applied. IFT20-mCherry and IFT54-mScarlet were excited using a 561 nm laser (Coherent), emission 575-625 nm. IFT27-GFP, Venus and G-GECO were excited using a 488 nm laser (Coherent), emission 500-550 nm. For two-channel imaging, simultaneous TIRF excitation was performed at 488 and 561 nm and a dual-view beam-splitter device was used to detect emission between 505-535 and 605-655 nm. Images were captured using NIS-Elements software (v3.1) at 100 ms per frame unless otherwise stated.

Fluorescence emission from single wavelength indicators such as G-GECO can be influenced by imaging artefacts such as flagella movement in the TIRF field of illumination. The use of a single emission wavelength calcium indicator was necessary for simultaneous imaging of IFT processes. We therefore performed a series of validation experiments to confirm that G-GECO was a reliable reporter of flagella Ca^2+^ elevations. 1) Rapid transient elevations in G-GECO fluorescence were not seen in the absence of external Ca^2+^ (Fig 3D). 2) G-GECO has a high dynamic range, exhibiting large changes in fluorescence emission on Ca^2+^ binding. Similar elevations in fluorescence were never observed in cells expressing CAH6-Venus, even during gliding motility (Supplementary Fig 1), indicating that motion artefacts do not give rise to large increases in fluorescence. 3) The nature of the [Ca^2+^]_fla_ elevations (e.g. duration, frequency) observed using G-GECO were very similar to those observed using ratiometric imaging of the fluorescent dyes Oregon Green BAPTA dextran and Texas Red dextran ^20^. Key findings using G-GECO were also verified using Oregon Green BAPTA dextran (e.g. Supplementary Fig 3B). 4) For detailed examination of [Ca^2+^]_fla_ and IFT, imaging of G-GECO was performed primarily in flagella immobilised with 0.1% poly-lysine to ensure that motion artefacts were minimised. 5) Dual channel imaging of G-GECO and IFT54-mScarlet enabled visual confirmation that changes in the intensity of G-GECO fluorescence were not replicated in the IFT54-mScarlet channel and were therefore due to changes in [Ca^2+^]_fla_ (Supplementary Fig 5C). This was particularly important for confirming that localised [Ca^2+^]_fla_ elevations were not artefacts due to flagella movement (e.g. Fig 4A, 5A).

Red fluorescent carboxylate-modified microspheres (0.1 μm diameter, Thermo) were used to image flagellar surface motility. Microspheres diluted in HEPES/NMG buffer were added to adherent cells and imaged by TIRF microscopy. Excitation with a 488nm laser alone was sufficient to simultaneously image G-GECO and microspheres due to the very high fluorescence emission of the microspheres. A dual-view beam-splitter device was used to detect emission between 505-535 (G-GECO or IFT27-GFP) and 605-655 nm (microspheres). Flagella exhibiting microsphere movements greater than 1 µm were examined for the presence of [Ca^2+^]_fla_ elevations during the course of microsphere movement.

For modulation of flagella adhesion with blue light, IFT54-MS cells were imaged by TIRF microscopy as previously, but a 488 nm laser was used to provide stimulation with blue light. Blue light was applied for 2 minutes to allow flagella to attach to the glass surface and then flagella lifting was examined after removing blue light for 30 seconds (defined as the removal of flagella from the surface, or a >50% decrease in flagella area in contact with the surface).

For imaging of flagella membrane glycoproteins, cells were stained with the fluorescent lectin concanavalin A (ConA) (100 μg mL^-1^) conjugated to fluorescein isothiocyanate (FITC) (Invitrogen, United Kingdom) for 5 minutes. Cells were then centrifuged (400 *g*, 1 minute) and resuspended in HEPES/NMG buffer prior to TIRF imaging.

### Data processing

Images were smoothed using a 3×3 pixel filter (NIS elements) and kymographs were generated using ImageJ software (version 1.45s; http://rsb.info.nih.gov/ij/). To identity [Ca^2+^]_fla_ elevations, we calculated the mean pixel intensity of G-GECO within a region of interest encompassing the whole flagellum, unless stated otherwise. Distal and proximal regions of interest were defined as regions 1 μm in length at either end of the flagellum. For quantitative analysis, the baseline intensity was calculated using an asymmetric lest squares smoothing algorithm (Origin) and changes in fluorescence emission were calculated using the ratio of fluorescence to the calculated baseline. Peaks were detected using Origin Pro software (peak finding method: local maximum with 5 local points), using a threshold value that differed from the baseline by 20%. IFT was monitored using TIRFM of fluorescently tagged IFT proteins. A kymograph of 60 s duration was used determine the mean velocity of IFT particles and the frequency at which they were observed within this time period. Entry of anterograde IFT trains was defined as the time at which the IFT train was first detected, as the region of the flagellum closest to the cell body may not be illuminated by TIRF excitation. IFT train velocity and frequency were quantified from the kymographs by measuring the slope and number of traces, respectively using ICY software ^67^. The interaction between IFT events and [Ca^2+^]_fla_ was examined by measuring relative G-GECO fluorescence coinciding with each event. To identify IFT events coinciding with the onset of [Ca^2+^]_fla_ elevations, a derivative Ca^2+^ trace was calculated by dividing the fluorescence in each frame by the median of the three preceding frames. The onset of each [Ca^2+^]_fla_ elevation was defined as the frame which displayed the greatest rate of change in fluorescence, and coinciding IFT events were defined as those events occurring within 0.5 s of this rise.

### Fluorescence recovery after photobleaching (FRAP)

For FRAP analysis of IFT, IFT54-MS cells were observed with a Nikon Eclipse Ti E inverted fluorescence microscope, equipped with a dual View DV2 (Photometrics) to detect emission between 505-535 and 605-655 nm, a iLAS FRAP module (Roper Scientifics) and Hamamatsu Flash 04 LT camera. Images were acquired using Metamorph software (version 7.7, Molecular Devices). IFT54-MS cells were placed in a poly-L-lysine (0.01%) treated dish and kept at room temperature during the acquisition. A segment of a flagellum was bleached by irradiation with a 150 mW green laser (Vortran), delivered through the 100X TIRF oil objective lens (100x, APO TIRF oil immersion, NA 1,49, Nikon) used for observation and focused on the specimen. The position to be photobleached was set at 50-70% of the flagellar length from the tip. The intensity of laser beam was 30%. Laser light for photobleaching was usually irradiated for about 1 sec. Images were recorded continuously for 15 sec before bleaching and then recovery was monitored for 45 sec by continuously recording videos with a 100 ms exposure.

### Transmission Electron Microscopy (TEM)

For chemical fixation of adherent *Chlamydomonas* flagella, cells were allowed to settle directly onto 0.01% poly-L-lysine coated coverslips for 5 min. The excess of cells was removed and cells were fixed with 1 % glutaraldehyde in 0.1 M sodium cacodylate (pH7.2) buffer for 10 min. Coverslips were washed 3 x 5 min in 0.1 M cacodylate buffer and then stained with 1 % uranyl acetate in distilled water for 1 h at 4°C. Coverslips were washed 3 x 10 min in distilled water. After serial dehydration with ethanol solution (30 %, 50 %, 70 %, 90 % et 3 x 100 %), samples were embedded in Agar 100 resin (Agar Scientific). Coverslips were mounted in Agar 100 capsule and left to polymerize at 60C for 2 days. To detach the coverslip, the sample was plunged into liquid nitrogen. First 200 nm ultrathin sections (50-70 nm) were collected in nickel grids Leica EM Uc 7 Ultramicrotome and stained with uranyl acetate (1 %) and lead citrate (80 mM). Observations were made on a Jeol 1400 TEM electron microscope. Images were processed with ImageJ.

## Acknowledgements

We thank Karl Lechtreck (Georgia), Joel Rosenbaum (Yale) and Luke Mackinder (York) for the generous supply of *Chlamydomonas* strains. The work was supported by a BBSRC award to GW (BB/M02508X/1) and an ERC Advanced Grant to CB (ERC-ADG-670390).

