## Supplementary data for "Flagella Ca^2+^ elevations regulate pausing of retrograde intraflagellar transport trains in adherent *Chlamydomonas* flagella"

**Supplementary Information**

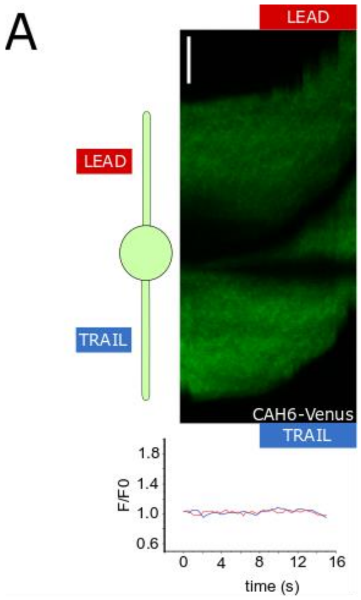


**Supplementary Fig 1: CAH6-Venus does not show elevations in fluorescence. A**) Kymograph showing gliding movements in cell expressing CAH6-Venus. The graph indicates that no significant changes in fluorescence occur in association with gliding movements (red =lead flagellum, blue = trailing flagellum). Bar = 5 μm.

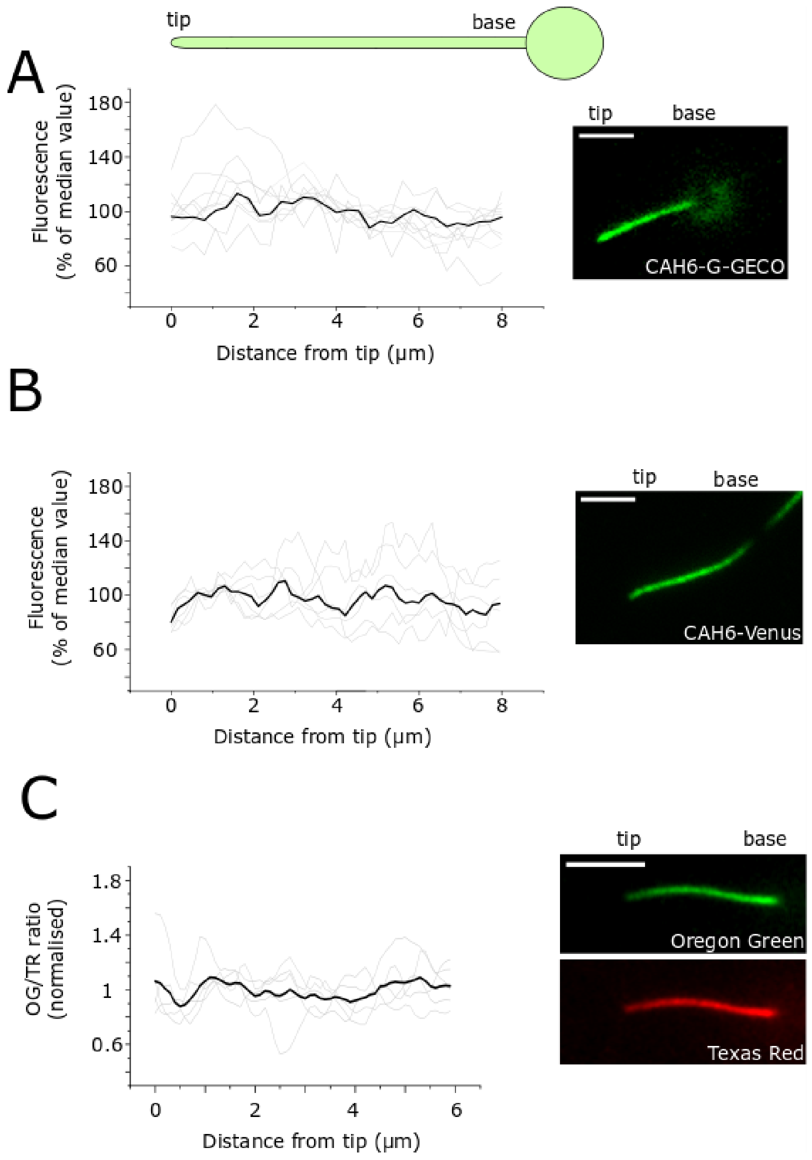


**Supplementary Figure 2: Resting $[Ca^{2+}]_{fla}$ is uniform along the length of the flagellum. A)** Median fluorescence of CAH6-G-GECO along the length of the flagellum. Individual traces (shown in grey) were normalised to the median fluorescence along the flagellum. n = 9 flagella. A representative TIRF microscopy image of flagellum expressing CAH6-G-GECO is shown. **B)** Median relative fluorescence of CAH6-Venus along the length of the flagellum. Individual traces (shown in grey) were normalised to the median fluorescence along the flagellum. n = 6 flagella. A representative TIRF microscopy image of flagellum expressing CAH6-Venus is shown. **C**) Median fluorescence ratio of Oregon green BAPTA and Texas Red along the length of the flagellum. Individual traces (shown in grey) were normalised to the median fluorescence ratio along the flagellum. n = 5 flagella. TIRF microscopy images of flagellum from a cell biolistically-loaded with Oregon-Green BAPTA dextran (green) and Texas Red dextran (red). Bars = 5 µm.

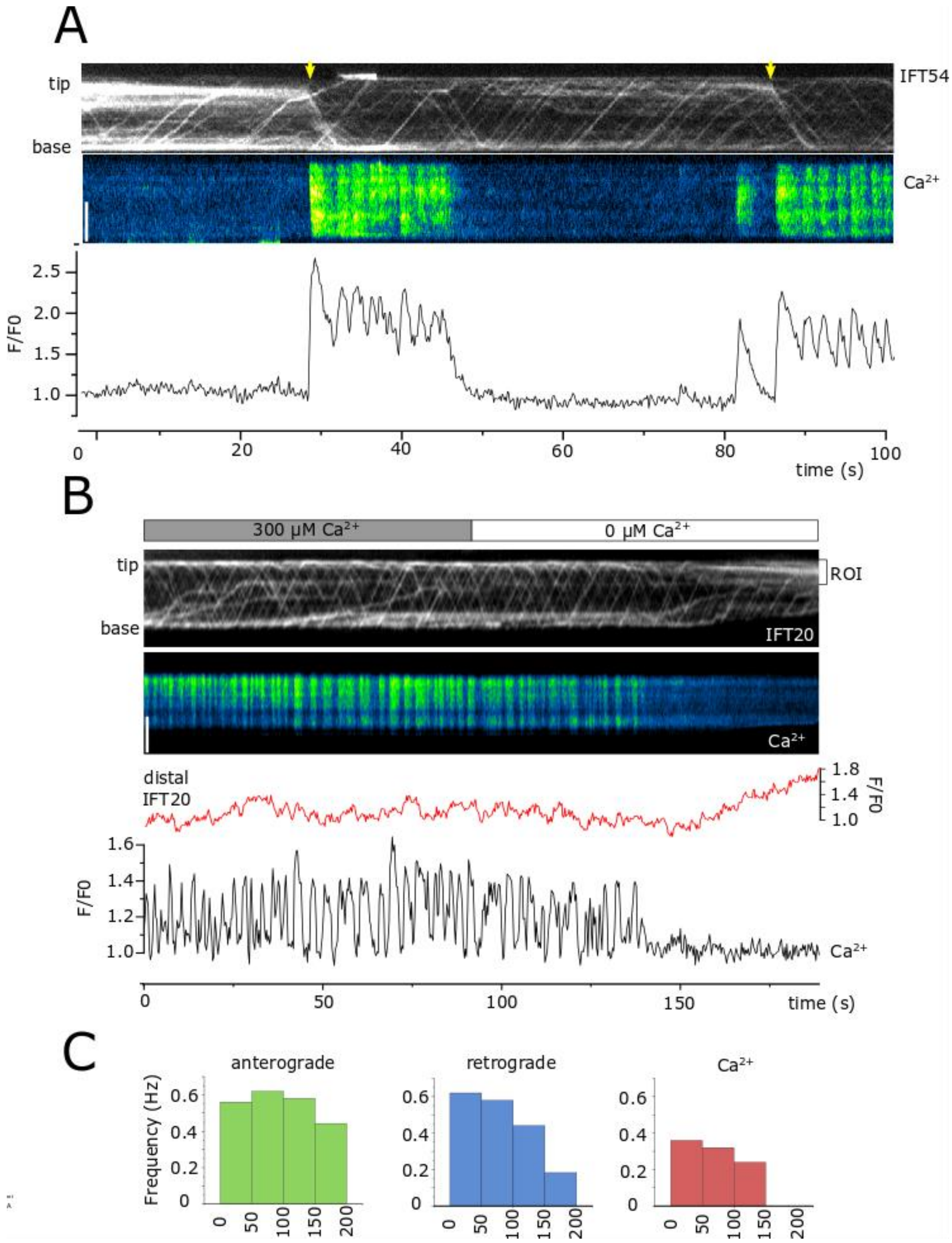


**Supplementary Figure 3: High frequency $[Ca^{2+}]_{fla}$ elevations prevent distal IFT accumulations. A)** Kymograph displaying simultaneous imaging of IFT54-mScarlet and G-GECO in a flagellum exhibiting intermittent $[Ca^{2+}]_{fla}$ elevations. In the absence of $[Ca^{2+}]_{fla}$ elevations, retrograde IFT trains accumulate in distal regions, but are cleared with the onset of $[Ca^{2+}]_{fla}$ elevations. Bar = 5 µm. **B)** Time course exhibiting the effect of removing external $Ca^{2+}$ on $[Ca^{2+}]_{fla}$ and IFT. The kymograph displays simultaneous imaging of IFT20-mCherry and Oregon Green BAPTA in a flagellum exhibiting repetitive $[Ca^{2+}]_{fla}$ elevations. The cell was initially perfused with a buffer containing 300 µM external $Ca^{2+}$, which was switched to a buffer containing 0 uM $Ca^{2+}$ after 100 s. The absence of external $Ca^{2+}$ inhibits $[Ca^{2+}]_{fla}$ elevations and leads to accumulation of IFT trains in the distal region of the flagellum (demonstrated by an increase of ITF20-mCherry fluorescence in the distal region - red trace). Note that pausing of anterograde IFT trains can occur during repetitive $[Ca^{2+}]_{fla}$ elevations. Bar = 5 µm. **C)** Bar charts indicating the frequency of $[Ca^{2+}]_{fla}$ elevations, anterograde IFT and retrograde IFT throughout the time course shown in (B).

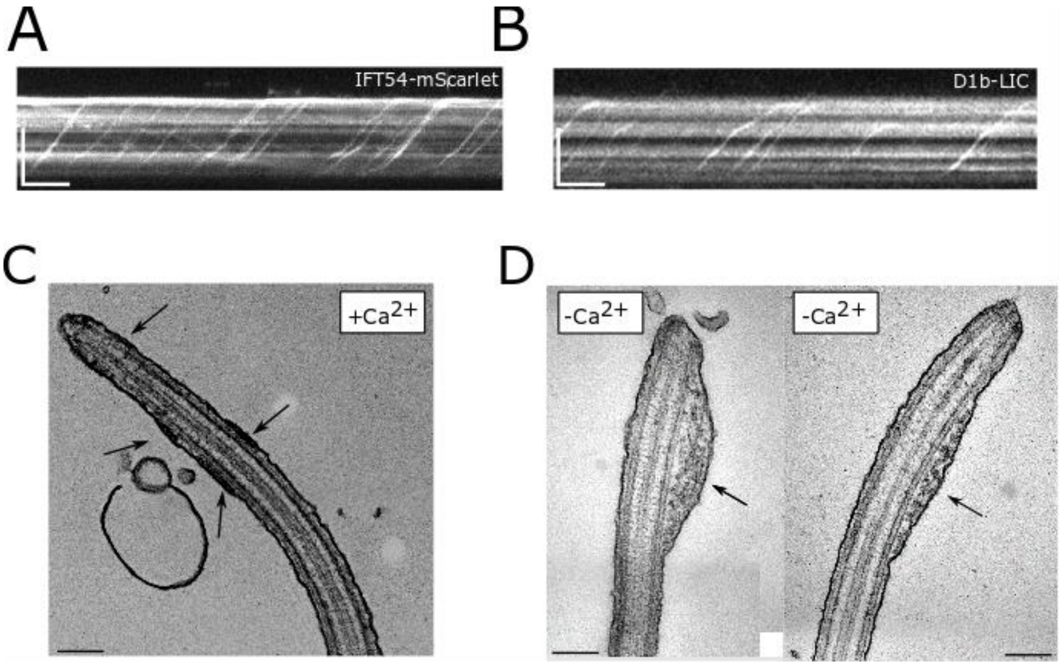


**Supplementary Figure 4: Inhibiting flagella $Ca^{2+}$ signalling causes large distal accumulations of IFT trains. A)** Kymograph of IFT54-mScarlet after removal of external $Ca^{2+}$ (0 $Ca^{2+}$ buffer + 200 uM EGTA) for 5 minutes. **B**) Kymograph of D1bLIC-GFP after removal of external $Ca^{2+}$ for 5 minutes. Bars = 5 µm and 10 s. **C**) TEM of adherent *Chlamydomonas* flagellum. Transverse section of flagellum indicating the presence of distinct IFT trains (arrowed). Bar = 300 nm. **D**) TEM of transverse sections of adherent flagella following the removal of external $Ca^{2+}$ for 5 minutes. Distinct IFT trains were not observed, but large accumulations of IFT particles can be observed in the distal regions of the flagella (arrowed). Bar = 300 nm.

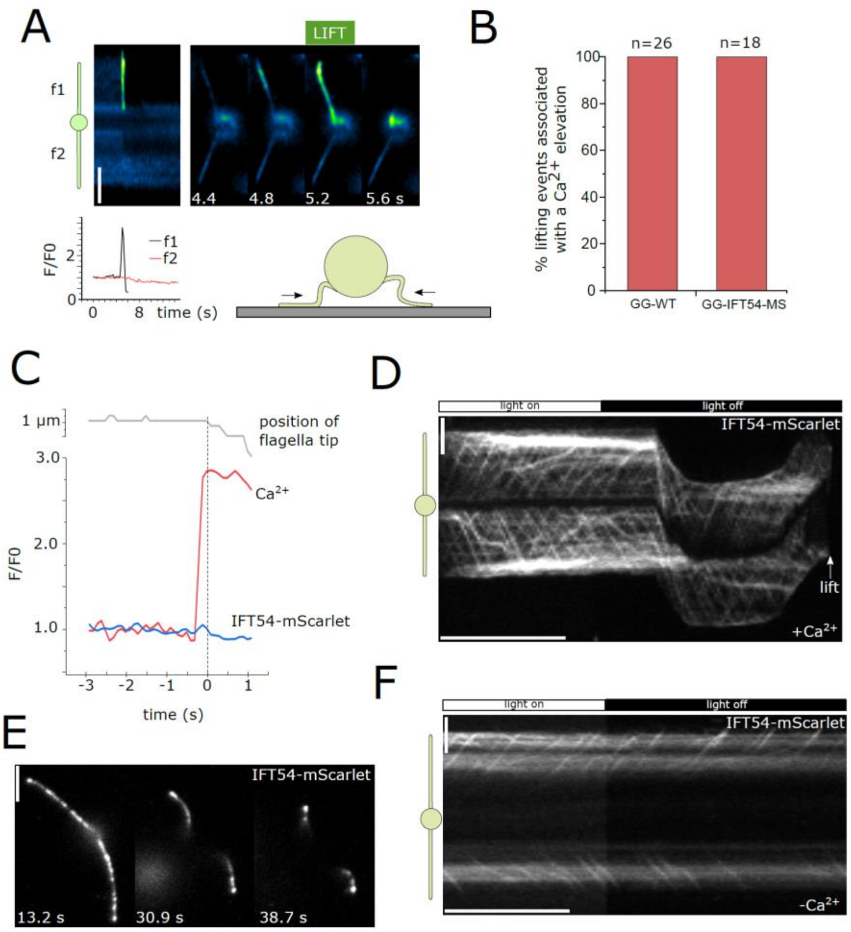


**Supplementary Figure 5: Flagella lifting is $Ca^{2+}$-dependent. A**) Imaging of $[Ca^{2+}]_{fla}$ elevations during flagella lifting in a GG-WT cell. The kymograph shows that a $[Ca^{2+}]_{fla}$ elevation directly precedes flagella lifting in the uppermost flagellum (f1). The trace below indicates that the $[Ca^{2+}]_{fla}$ elevation is restricted to the lifting flagellum only. Bar = 5 µm. **B**) Quantitation of $[Ca^{2+}]_{fla}$ elevations associated with flagella lifting in GG-WT and GG-IFT54-MS cells. The percentage of lifting events that coincide with a $[Ca^{2+}]_{fla}$ elevation is shown. **C)** Detailed examination of the timing of $Ca^{2+}$ signalling during flagella lifting in GG-IFT54-MS cells. The $[Ca^{2+}]_{fla}$ elevation (red) precedes the movement of the flagellum (shown by the position of the flagella tip – grey line) by 200 ms. Note that the increase in G-GECO fluorescence was not accompanied by any significant increase in the fluorescence of IFT54-mScarlet (blue line) indicating that changes in G-GECO fluorescence were not the result of movement artefacts. **D)** Kymograph indicating the response of a IFT54-MS cell to the removal of blue light. Movement and subsequent lifting of flagella caused by the removal of blue light (488 nm) is associated with removal of accumulations of paused retrograde IFT trains in the distal region. Bars = 5 µm and 15 s. **E**) Image series showing the response of IFT54-MS cell to the removal of blue light. The cells adopts the typical gliding configuration, with flagella arranged at 180° to each other, in the presence of blue light. . Removal of blue light promotes withdrawal of flagella towards the cell body, so that only the flagella tips remain adherent. Bar = 5 µm. **F**) In the absence of external $Ca^{2+}$, the removal of blue light does not have any impact on accumulated IFT trains and flagella lifting did not occur. Bars = 5 µm and 15 s.

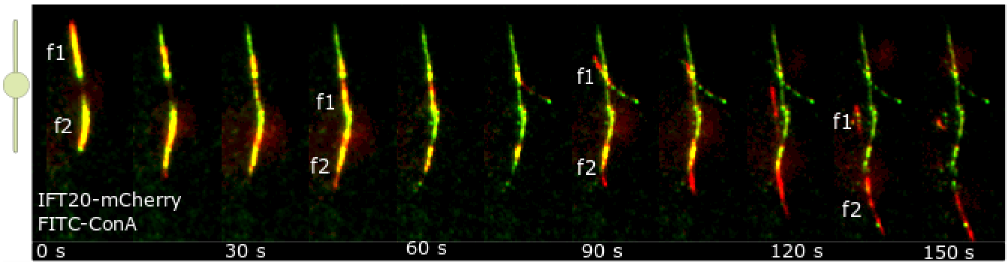


**Supplementary Figure 6: Labelling of FMG-1B with fluorescent lectin.** Image series of gliding IFT20-mCherry (red) cell viewed by TIRF microscopy. The cell was labelled with the fluorescent lectin, fluorescein isothiocyanate- concanavalin A (FITC-ConA, green) prior to being allowed to settle. FITC-ConA labels the major adhesive glycoprotein, FMG-1B, in the flagella membrane. FMG-1B is shed from flagella during gliding and lifting movements, leaving behind a trail of the adhesive glycoprotein.

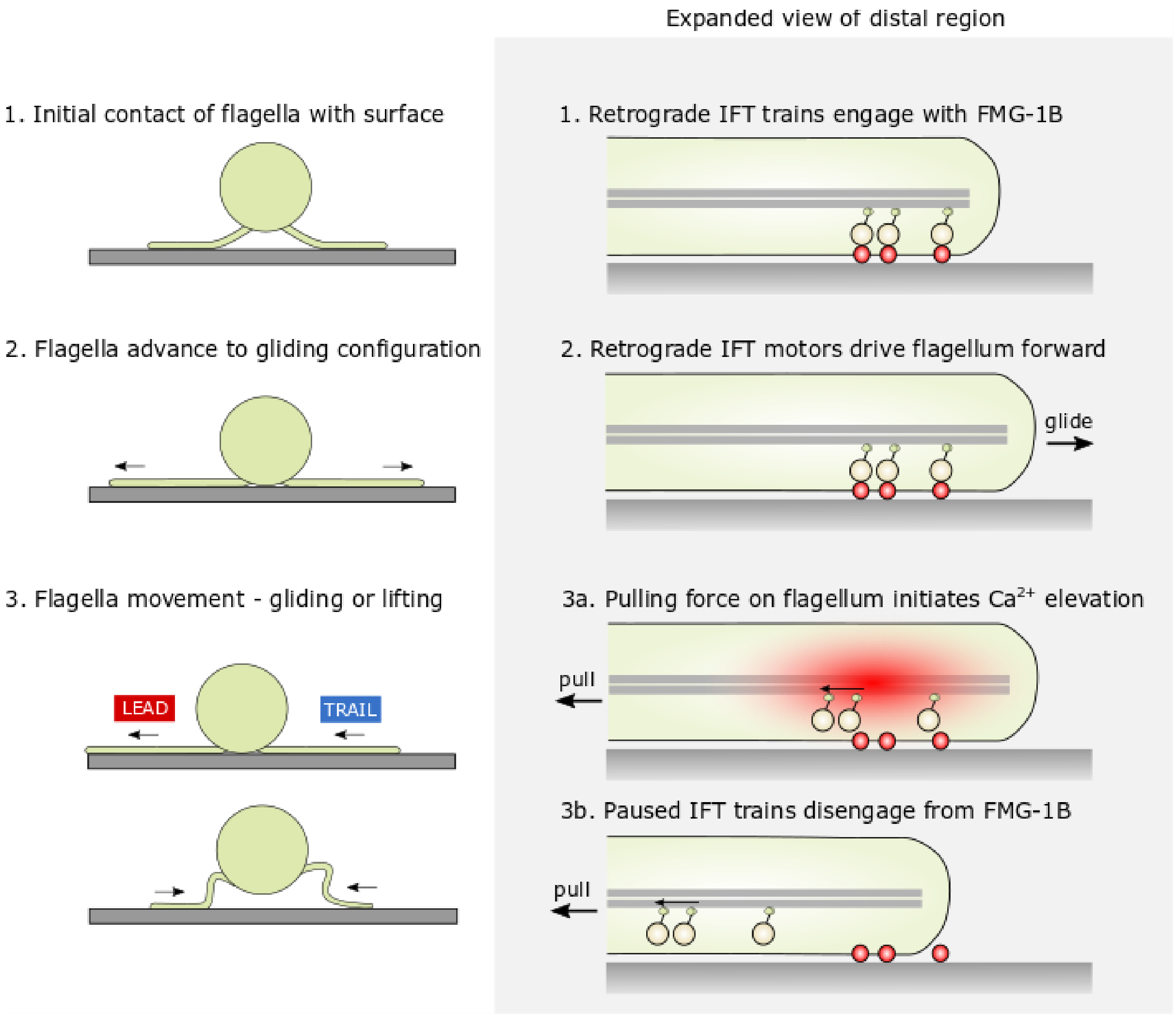


**Supplementary Figure 7: Proposed scheme of the interaction between $Ca^{2+}$ and retrograde IFT trains.** 1) In a gliding flagellum, retrograde IFT trains interact with adherent FMG-1B in the flagella membrane. 2) The accumulation of paused retrograde IFT trains pulls the flagellum forward (gliding motility) until the flagella are fully extended. The accumulation of paused retrograde IFT trains bound to FMG-1B means that flagellum resists pulling forces from the opposite direction (towards the cell body), caused either by bending of the axoneme or by the other gliding flagellum. 3a) If sufficient pulling force is applied, a mechanosensitive $Ca^{2+}$ elevation occurs in the flagellum that disrupts the interaction between FMG-1B and the paused retrograde IFT trains, causing the retrograde IFT trains to return to the cell body. 3b) FMG-1B can therefore move freely in the flagella membrane, removing the resistance to the pulling force, and allowing the flagellum to be dragged along the surface.
